# A redox-regulated RCC1-like protein controls catalase activity in Arabidopsis

**DOI:** 10.64898/2026.01.30.702735

**Authors:** Zhoubo Hu, Sara Christina Stolze, Amna Mhamdi, Frank Van Breusegem, Hirofumi Nakagami, Roman Ulm

## Abstract

Reactive oxygen species (ROS) are central regulators of plant growth and stress responses. Cellular ROS levels are tightly controlled by antioxidant systems, including the evolutionarily conserved catalases that detoxify hydrogen peroxide (H_2_O_2_) predominantly within peroxisomes. Despite their importance, substantial gaps remain in our understanding of catalase biogenesis, regulation, subcellular targeting, and potential extra-peroxisomal functions. Using affinity purification of the UV-B photoreceptor UVR8 coupled with mass spectrometry, we identified a REGULATOR OF CHROMATIN CONDENSATION 1–like protein in *Arabidopsis*, which we named CATALASE-INTERACTING RCC1-LIKE 1 (CAIR1). CAIR1 interacts with all three catalase isoforms (CAT1–CAT3) as well as their chaperone NO CATALASE ACTIVITY 1 (NCA1). Loss-of-function *cair1* mutants partially phenocopy *cat2* and *nca1*, with reduced catalase activity, enhanced sensitivity to oxidative stress and alkaline growth conditions, and impaired primary root elongation. Mechanistically, cytosolic interaction between CAIR1 and CAT2 enhances total cellular catalase activity by facilitating peroxisomal import and proper subcellular localization of CAT2. In the absence of CAIR1, CAT2 forms aggregates, likely accounting for the observed loss of catalase activity. Notably, CAIR1 undergoes reversible, redox-dependent oligomerization that enhances its interaction with catalases. Mutation of CAIR1 at Cys-356 and Cys-545 compromises this interaction under elevated ROS conditions and fails to rescue the oxidative stress sensitivity of *cair1* mutants. Moreover, UV-B exposure suppresses catalase activity by weakening the interaction between CAIR1 and catalases, thus linking environmental light signalling to cellular redox regulation. Together, our findings reveal CAIR1 as a dynamic redox-responsive regulator of catalase activity that maintains cellular redox homeostasis by coordinating catalase localization and function through reversible oligomerization.

## Introduction

The build-up of reactive oxygen species (ROS) within various cellular compartments in response to stress can lead to oxidative stress, ultimately compromising both cellular and organellar integrity. Key ROS include singlet oxygen (^1^O_2_), superoxide anion (O_2_^•−^), hydrogen peroxide (H_2_O_2_), and hydroxyl radicals (•OH); species that can interconvert within the cellular environment. Despite their damaging potential, ROS also serve as important signalling molecules that initiate pathways involved in plant growth, development, and adaptation to stress (Apel & Hirt, 2004; Chi *et al*, 2013; Mittler, 2017; Mittler *et al*, 2022). As such, maintaining a delicate balance in ROS levels is essential. For ROS homeostasis, plants rely on a complex network of non-enzymatic antioxidants and processing enzymes such as glutathione, ascorbate, ascorbate peroxidases, superoxide dismutases, and catalases (CAT), all of which are crucial for sustaining ROS homeostasis (Mhamdi *et al*, 2010; Mittler *et al*, 2022).

Peroxisomes are single-membrane organelles involved in a variety of metabolic processes. In plants, peroxisomes play essential roles in key metabolic pathways such as photorespiration and lipid mobilization through fatty acid β-oxidation (Del Rio & Lopez-Huertas, 2016; Hu *et al*, 2012; Kao *et al*, 2018; Mhamdi *et al*, 2012; Molina-Moya *et al*, 2025; Pan *et al*, 2020). Photorespiration is a metabolic process in plants whereby the enzyme ribulose-1,5-bisphosphate carboxylase/oxygenase (Rubisco) fixes oxygen instead of carbon dioxide, resulting in reduced photosynthetic efficiency and the production of harmful ROS in peroxisomes (Foyer *et al*, 2009). Fatty acid β-oxidation provides energy for post-germinative growth and development by breaking down fats stored as triacylglycerol in oil bodies of oilseed plants such as *Arabidopsis thaliana* (herein Arabidopsis) (Theodoulou & Eastmond, 2012). Both fatty acid β-oxidation and photorespiration generate H_2_O_2_ in peroxisomes, which is predominantly scavenged by catalases (Baker *et al*, 2023; Tuzet *et al*, 2019). Catalases are tetrameric haem-containing antioxidant enzymes that catalyze the breakdown of H_2_O_2_, and thereby play a critical role in maintaining H_2_O_2_ homeostasis (Kamigaki *et al*, 2003; Kato *et al*, 2021; Reumann *et al*, 2004; Scandalios *et al*, 1980). Import of proteins such as catalases into peroxisomes for these essential functions primarily depends on peroxisome targeting signals (PTS) (Cross *et al*, 2016). Proteins with PTS require cytosolic PEROXIN-5 (PEX5) and PEX7 proteins for import (Gould *et al*, 1989; Kato *et al*, 1996; Nito *et al*, 2002), whereas PTS-independent access to peroxisomes is regulated by “piggy-back” co-import with a peroxisomal-targeted, PTS-containing protein (Glover *et al*, 1994; Lee *et al*, 1997). Outside of peroxisomes, catalases are localized to and perform roles in the nucleus and cytosol, as recently described (Al-Hajaya *et al*, 2022; Baker *et al*, 2023; He *et al*, 2021; Lin *et al*, 2025). Indeed, mammalian catalases play key roles in defending against acute oxidative stress in the cytosol (Dubreuil *et al*, 2020) and cytosolic catalase levels are regulated directly at the level of their peroxisomal import (Okumoto *et al*, 2020).

As in most flowering plants, the Arabidopsis genome encodes three catalase isoforms, namely CAT1, −2 and, −3 (Frugoli *et al*, 1996; Mhamdi *et al*, 2010). Among these, CAT2 is the major isoform in leaves, accounting for approximately 90% of total catalase activity (Mhamdi *et al*, 2012; Mhamdi *et al*, 2010; Su *et al*, 2018; Yang *et al*, 2019). Arabidopsis catalases are imported into peroxisomes by a non-canonical import pathway (Baker *et al*, 2023; Oshima *et al*, 2008). Although the C-terminal 11 amino acids were shown to be required for plant catalase localization to peroxisomes in transient expression systems (Fujikawa *et al*, 2019), they are not essential for peroxisomal targeting in stably transformed Arabidopsis plants (Al-Hajaya *et al*, 2022; Kamigaki *et al*, 2003). Furthermore, different interactors of catalases were identified and linked to the regulation of catalase activity, including calcium-regulated calmodulin (CaM), the zinc finger protein LESION SIMULATING DISEASE 1 (LSD1), NUCLEOREDOXIN 1 (NRX1), and NO CATALASE ACTIVITY 1 (NCA1) as positive regulators (Baker *et al*, 2023; Kneeshaw *et al*, 2017; Li *et al*, 2013; Lin *et al*, 2025; Lv *et al*, 2019; Yang & Poovaiah, 2002). Notably, NCA1, a protein with an N-terminal RING-finger domain and a C-terminal tetratricopeptide repeat-like helical domain, interacts with catalases in the cytosol, where it chaperones catalases to maintain stability and activity, hence playing an important role in multiple stress responses, including in response to enhanced photorespiration (Hackenberg *et al*, 2013; Li *et al*, 2015; Liu *et al*, 2019).

Mammalian and fungal Regulator of Chromatin Condensation 1 (RCC1) proteins are conserved eukaryotic chromatin-associated proteins consisting of seven RCC1-repeats that function as guanine nucleotide exchange factors (GEFs) for the small GTPase Ran. However, several proteins with one or more RCC1-like structural domains (RLDs) of 51–68 amino acid RCC1-like repeats exhibit diverse non-GEF functions (Hadjebi *et al*, 2008). The Arabidopsis genome encodes 24 RLD-containing proteins (Tilbrook *et al*, 2013), of which only a few are functionally characterized and shown to be involved in a diverse range of functions, including efficient splicing of *nad2* mRNA and accumulation of mitochondrial respiratory chain complex I (Kühn *et al*, 2011; Su *et al*, 2017), plasticity of rosette size in response to nitrogen availability (Duarte *et al*, 2021), abscisic acid signalling (Ji *et al*, 2019), cold acclimation and freezing tolerance (Ji *et al*, 2015), gravity signalling and regulation of polar auxin transport (Furutani *et al*, 2020), and ultraviolet-B (UV-B) sensing and signalling (Podolec *et al*, 2021; Rizzini *et al*, 2011). The UV-B photoreceptor UV RESISTANCE LOCUS 8 (UVR8) is a member of the RCC1-like protein family in plants (Kliebenstein *et al*, 2002; Rizzini *et al*, 2011) that, however, has no Ran-GEF activity (Brown *et al*, 2005) and does not directly bind to chromatin (Binkert *et al*, 2016). The cytosolic homodimeric form of UVR8 monomerizes upon UV-B photon reception, after which it accumulates in the nucleus to regulate downstream gene expression, thus mediating UV-B dependent photomorphogenic responses including acclimation and UV-B stress protection (Podolec *et al*, 2021).

Here, we identify a previously uncharacterized RCC1-like protein, CATALASE-INTERACTING RCC1-LIKE 1 (CAIR1), as a regulator of catalase activity in Arabidopsis. CAIR1 associates with all three catalases and their chaperone NCA1 in the cytosol and is required for normal catalase function in vivo. Our results indicate that CAIR1 function is modulated by the cellular redox state, uncovering a regulatory link between catalase homeostasis and dynamic redox signalling.

## Results

### UVR8 co-purifies and interacts with AT5G11580, an RCC1-like family protein of unknown function

We generated an N-terminally GFP-tagged UVR8 genomic clone by fast-track BAC recombineering (Hu *et al*, 2019) that complemented the *uvr8-6* null mutant allele (Supplemental Figure S1A–C). We used this line for AP–MS using anti-GFP nanobodies/VHH coupled to agarose beads (GFP-trap®) for complex purification, which identified previously known UVR8 core photocycle proteins REPRESSOR OF UV-B PHOTOMORPHOGENESIS 1 (RUP1), RUP2, CONSTITUTIVELY PHOTOMORPHOGENIC 1 (COP1), as well as the COP1-associated SUPPRESSOR OF PHYA-105 (SPA) proteins SPA1-to-SPA4, specifically when plants were grown in the presence of supplementary UV-B (Supplemental Figure S1D and Supplemental Table S1) (Favory *et al*, 2009; Gruber *et al*, 2010; Heijde *et al*, 2013). In addition, a RCC1-like protein (AT5G11580; herein CAIR1, see below) of previously unknown function co-purified, both in the absence and presence of UV-B (Supplemental Figure S1D). Consistently, CAIR1 interacted with UVR8 in co-immunoprecipitation (co-IP) assays when co-expressed in human embryonic kidney (HEK293) cells, as well as in luciferase complementation imaging (LCI) assays in *Nicotiana benthamiana* epidermal leaf cells (Supplemental Figure S1E, F).

CAIR1 is a 553-amino protein that contains six predicted tandem RCC1 repeats but no additional predicted functional domain (Supplemental Figure S2). To investigate the potential role of CAIR1 in UVR8 signalling, we isolated the *cair1-1* mutant containing a T-DNA insertion in the second intron and generated the additional *cair1-2* and *cair1-3* null alleles by CRISPR/Cas9 (Supplemental Figure S3). Loss and overexpression of CAIR1 did not obviously affect well-known UVR8–mediated UV-B responses such as hypocotyl growth inhibition, anthocyanin accumulation, accumulation of UV-absorbing metabolites, activation of UV-B–responsive marker genes (*RUP2*, *HY5*, and *CHS*), and UV-B acclimation (Supplemental Figure S4). We thus identify CAIR1 as a previously undescribed interactor of the UVR8 photoreceptor, with our findings indicating that its functional relevance may extend beyond the canonical UVR8 signalling pathway underlying UV-B–induced photomorphogenesis.

### CAIR1 (AT5G11580) interacts with catalases and their chaperone NCA1

To gain insight into the molecular function of CAIR1, we generated Pro_35S_:YFP-CAIR1 and Pro_35S_:CAIR1-YFP lines in the respective *cair1-1* null mutant background, as well as a genomic complementation line containing a C-terminal GFP tag. AP–MS was performed using GFP-trap agarose beads on proteins extracted from 15-d-old transgenic plants. As controls, protein extracts from wild-type and Pro_35S_:StrepII-3xHA-YFP (herein “YFP”) seedlings were processed in parallel to account for potential common contaminant (non-specifically binding) proteins. CAIR1-GFP expressed from the genomic complementation construct indeed reciprocally co-purified UVR8, but in all three cases (Pro_35S_:YFP-CAIR1, Pro_35S_:CAIR1-YFP, as well as genomic Pro_CAIR1_:CAIR1-GFP), tagged CAIR1 powerfully co-purified the three catalases CAT1, CAT2, and CAT3, as well as NCA1 (Figure 1A, and Supplemental Table S1); hence the name CATALASE INTERACTING RCC1-LIKE 1 (CAIR1) assigned for the previously undescribed AT5G11580. In agreement with the AP–MS data, CAIR1-GFP efficiently co-immunoprecipitated endogenous catalases (Figure 1B), which were detected with an anti-catalase antibody that cross-reacts with all three catalases (Su *et al*, 2018). Moreover, immunoprecipitation of GFP-CAT2 and NCA1-GFP from functional genomic complementation lines *nca1-1/*Pro_NCA1_:NCA1-GFP and *cat2-2/*Pro_CAT2_:GFP-CAT2 (Supplemental Figure S5A–D), respectively, both co-immunoprecipitated endogenous CAIR1 (Figure 1C and 1D). Additionally, AP–MS of NCA1-GFP and GFP-CAT2 further corroborate that CAIR1 forms a complex with catalases and NCA1 (Supplemental Table S1; and see below). To complement the AP–MS and co-IP findings, we performed yeast two-hybrid (Y2H) assays to test for potential direct protein–protein interactions. Y2H assays confirmed that CAIR1 interacts with CAT1, CAT2, and CAT3, but notably did not show CAIR1 interaction with NCA1 (Figure 1E and Supplemental Figure S6A). However, interaction between NCA1 and CAT2 was detectable (Supplemental Figure S6A). In an additional orthogonal assay, heterologously expressed GFP-CAIR1 (but not GFP alone) in HEK293 cells co-immunoprecipitated co-expressed Flag-CAT2 and Flag-NCA1 (Supplemental Figure S6B, C). LCI assays in *N. benthamiana* epidermal leaf cells further indicated that CAIR1 interacts with both CAT2 and NCA1 (Figure 1F). By contrast, no interactions were identified in the LCI assays for negative control combinations of CAIR1 with RUP2, or CAT2 and NCA1 with either UVR8 or RUP2, although UVR8 and RUP2 exhibited strong interaction, thus supporting functional UVR8 and RUP2 expression (Figure 1F). Altogether, we conclude that Arabidopsis CAIR1 interacts and forms stable complexes with catalases and their chaperone NCA1.

**Figure 1.**
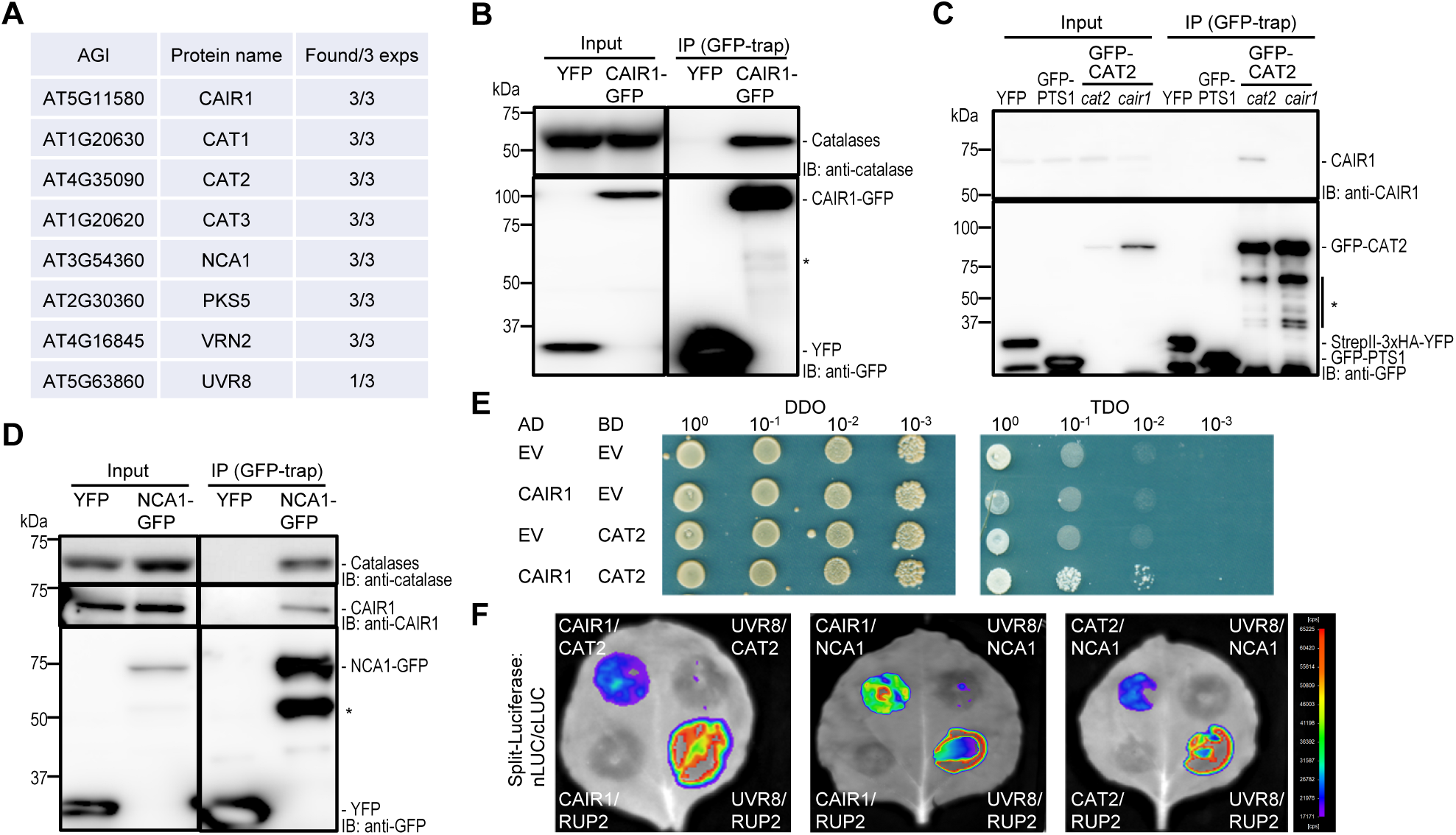
CAIR1 interacts with catalases and NCA1. **(A)** Summary of proteins identified in three independent AP–MS experiments of CAIR1 complexes (Supplementary Table S1). Depicted is a concise list of candidate proteins significantly enriched in all three CAIR1 AP–MS experiments compared to in the negative controls (wild type and Pro_35S_:StrepII-3xHA-YFP). Note that the 3 experiments represent AP–MS with *cair1-1*/Pro_35S_:YFP-CAIR1, *cair1-1*/Pro_35S_:CAIR1-YFP, and genomic complementation line *cair1-1*/Pro_CAIR1_:CAIR1-GFP; UVR8 co-purified with the latter. AGI = Arabidopsis Gene Identifier. **(B)** Co-IP assay of endogenous catalases using GFP-trap purification from extracts of 15-d-old Pro_35S_:StrepII-3xHA-YFP (YFP) and *cair1-1*/Pro_CAIR1_:CAIR1-GFP (CAIR1-GFP) transgenic seedlings. **(C)** Co-IP assay of endogenous CAIR1 using GFP-trap purification from extracts of 15-d-old Pro_35S_:StrepII-3xHA-YFP (YFP), Pro_35S_:GFP-PTS1 (GFP-PTS1), *cat2-2*/Pro_CAT2_:GFP-CAT2 (GFP-CAT2 *cat2*), and *cair1-1*/Pro_CAT2_:GFP-CAT2 (GFP-CAT2 *cair1*) transgenic seedlings. **(D)** Co-IP assay of endogenous CAIR1 and catalases using GFP-trap purification from extracts of 15-d-old Pro_35S_:StrepII-3xHA-YFP (YFP) and *nca1-1*/Pro_NCA1_:NCA1-GFP (NCA1-GFP) transgenic seedlings. (B–D) YFP and/or GFP-PTS1 transgenic lines were used as negative controls. IB, immunoblotting; IP, immunoprecipitation; asterisks (*) indicate likely degradation products that evolved during immunoprecipitation. **(E)** Interaction of CAIR1 with CAT2 in a yeast two-hybrid growth assay. Tenfold serial dilutions of transformed yeast spotted on SD/−Trp/−Leu (DDO; nonselective for interaction) and SD/−Trp/−Leu/−His (TDO; selective) plates. AD, activation domain; BD, DNA binding domain; EV, empty vector. **(F)** LCI assay using indicated combinations of 35S promoter–driven CAIR1-nLUC, cLUC-CAT2, cLUC-NCA1, UVR8-nLUC, and cLUC-RUP2 transient expression in *N. benthamiana* leaves. Results shown are representative of 3 independent repetitions.

### CAIR1 localizes to the cytosol, where it interacts with CAT2

In addition to showing that CAIR1 forms a complex which includes catalases and NCA1 (Figure 2A, Supplemental Table S1), both NCA1-GFP and GFP-CAT2 AP–MS provided a comprehensive insight into the CAT2 interactome. For AP–MS of GFP-CAT2, a peroxisomal marker line (Pro_35S_:GFP-PTS1) (Mano *et al*, 2002) was used as an additional negative control next to wild type and a YFP-expressing line to reliably and robustly identify CAT2 interactors, including those in the peroxisome. In total, 82 proteins were enriched in GFP-CAT2 purifications compared with wild type, YFP, and GFP-PTS1 controls, many of which have been experimentally shown to localize to peroxisomes or possess a putative PTS (Supplemental Figure S7A, Supplemental Table S1 and S3). Next to the top co-purifying proteins CAIR1, NCA1, CAT1, and CAT3, as based on fold-enrichment, several previously known CAT2 interactors were identified in the GFP-CAT2 AP–MS analysis, including PEX5 (Kamigaki *et al*, 2003; Williams *et al*, 2012), ACYL-COA OXIDASE 4 (ACX4) and ACX5 (Yuan *et al*, 2017), GLYCOLATE OXIDASE 1 (GOX1), GOX2 (Yang *et al*, 2025; Zhang *et al*, 2016), and *S*-NITROSOGLUTATHIONE REDUCTASE (GSNOR1) (Chen *et al*, 2020; Zhang *et al*, 2024) (Figure 2A). In addition, we identified the peroxisomal HYDROXYPYRUVATE REDUCTASE 1 (HPR1) as a potential, previously undocumented interactor of CAT2 (Figure 2A). While GFP-CAT2 co-immunoprecipitated endogenous HPR1 as well as CAIR1 (Supplemental Figure S7B), neither CAIR1-GFP nor NCA1-GFP pulled down HPR1, despite robust co-IP of catalases (Supplemental Figure S7C, D). These results are consistent with the AP–MS data and the subcellular localizations of these proteins, and they suggest that CAIR1 may interact specifically with non-peroxisomally localized catalases. It is moreover noteworthy that AP–MS of CAIR1-GFP co-immunoprecipitated CATs and UVR8, but UVR8 and CAT could not be found in GFP-CAT2 and GFP-UVR8 AP–MS, respectively, despite clear co-IP of CAIR1 in both cases (Figure 2A and Supplemental Figure S1D), suggesting the absence of a ternary UVR8-CAIR1-CAT protein complex.

**Figure 2.**
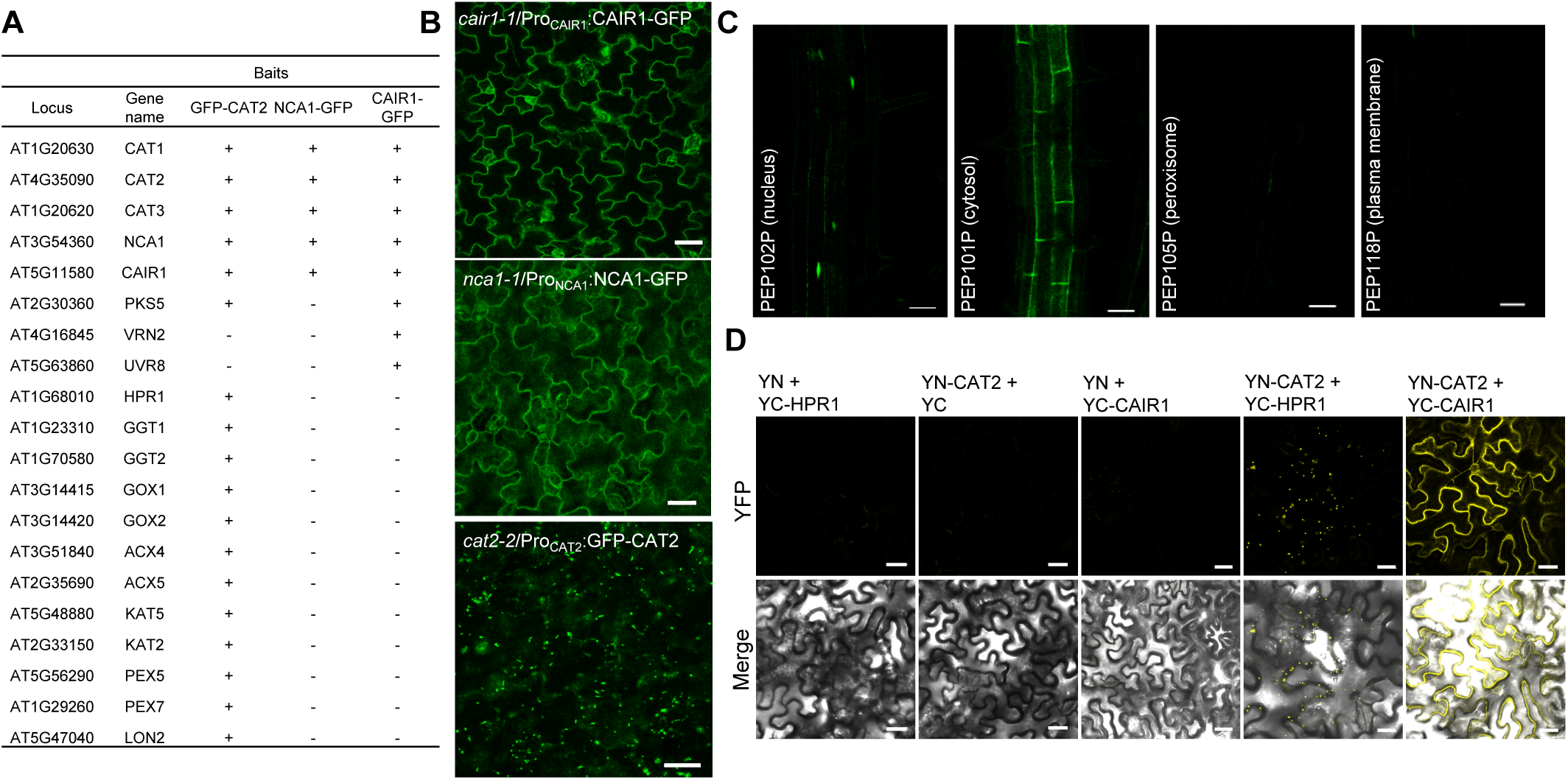
CAIR1 localizes to the cytosol where it interacts with CAT2. **(A)** List of proteins identified in the purified GFP-CAT2, NCA1-GFP, and CAIR1-GFP complexes. Representation of peptide peaks of different subunits measured by LC-MS/MS (Supplementary Table S1) is indicated schematically (+). **(B)** Subcellular localization of CAIR1-GFP, NCA1-GFP, and GFP-CAT2 in stably transformed 5-d-old Arabidopsis seedlings. Scale bars: 30 μm. **(C)** CAIR1 localizes to nuclei and cytosol but not to peroxisomes or plasma membrane. CAIR1 fused with sfGFP^11^ at the C-terminus was introduced into transgenic Arabidopsis expressing sfGFP1-10^OPT^ targeted to peroxisomes (PEP105P), nuclei (PEP102P), cytosol (PEP101P), and plasma membrane (PEP118P). GFP signals were examined using confocal laser scanning microscopy. Scale bar: 30 μm. **(D)** BIFC assay in *N. benthamiana* leaves transiently expressing YFP fusion proteins YFPN-CAT2 (YN-CAT2), YFPC-HPR1 (YC-HPR1), and YFPC-CAIR1 (YC-CAIR1). Depicted are YN-CAT2–YC-HPR1 interaction in peroxisomes, and YN-CAT2–YC-CAIR1 interaction in the cytosol. Scale bars: 30 μm.

Although catalase activity and localization have been found in various cellular compartments, including the cytosol, mitochondria, chloroplasts, and nucleus, catalases are predominantly localized in peroxisomes (Al-Hajaya *et al*, 2022; Baker *et al*, 2023; Mhamdi *et al*, 2012). Native promoter–driven CAIR1-GFP displayed cytosolic localization, mirroring the cytosolic localization of NCA1-GFP (Figure 2B). Despite the demonstrated interaction of CAT2 with both CAIR1 and NCA1, GFP-CAT2 exhibited primarily peroxisomal localization (Figure 2B). To further investigate the subcellular localization of CAIR1 with higher sensitivity, we used the self-assembling split superfolder GFP (sfGFP^OPT^) system (Al-Hajaya *et al*, 2022; Park *et al*, 2017). The co-expression of CAIR1-sfGFP11 with sfGFP1-10 β-strand (sfGFP1-10^OPT^) targeted to cytosol and nucleus resulted in reconstituted sfGFP fluorescence signals detectable by confocal laser scanning microscopy, which were not observed with peroxisome and plasma membrane targeted sfGFP1-10^OPT^ (Figure 2C), suggesting that CAIR1 is mainly localized to the cytosol and nucleus. It is of note that, in addition to NCA1 cytosolic and catalase peroxisomal localization, nuclear localization has also been described for both (Al-Hajaya *et al*, 2022; Hackenberg *et al*, 2013). Thus, to reveal the subcellular localization of the CAIR1–CAT2 interaction, we used the bimolecular fluorescent complementation (BiFC) assay (Kerppola, 2006) in transiently transformed *N. benthamiana* epidermal leaf cells. Reconstituted cytosolic YFP signals were detected when YC-CAIR1 and YN-CAT2 were co-expressed, indicating interaction of CAIR1 and CAT2 in the cytosol; however, YFP signal indicating interaction was not observed in nuclei or peroxisomes (Figure 2D). This specific cytosolic interaction was further supported by comparing it to the sharply contrasting localization of YN-CAT2–YC-HPR1 interaction detected only in peroxisomes (Figure 2D). We conclude that, whereas HPR1 identified in our GFP-CAT2 AP–MS is a previously undescribed interactor of CAT2 in peroxisomes, the CAIR1 interaction with CAT2 is — like the NCA1–CAT2 interaction — confined to the cytosol.

### *cair1* mutants show reduced catalase activity associated with catalase aggregation

Given the clear CAIR1–CAT interaction, we tested whether CAIR1 affects catalase activity. Indeed, catalase activities of 5-d-old *cair1-1, cair1-2*, and *cair1-3* seedlings were reduced when compared to that in wild type and two complementation lines (Figure 3A). In contrast to *nca1* (Hackenberg *et al*, 2013), the reduced catalase activities in the *cair1* mutant alleles were not associated with reduced catalase protein levels (Figure 3B, and Supplemental Figure S3C). Thus, to better understand the reason for reduced catalase activity, we further assessed whether CAIR1 affected CAT2 subcellular localization. Interestingly, GFP-CAT2 formed aggregates in the *cair1* background but not in *cat2* complementation lines (Figure 3C). In contrast, the peroxisome marker GFP-PTS1 showed no difference in subcellular localization when expressed in *cair1* versus in wild type (Figure 3C). We conclude that CAIR1 interaction with CAT2 in the cytosol ensures proper peroxisomal localization of CAT2.

**Figure 3.**
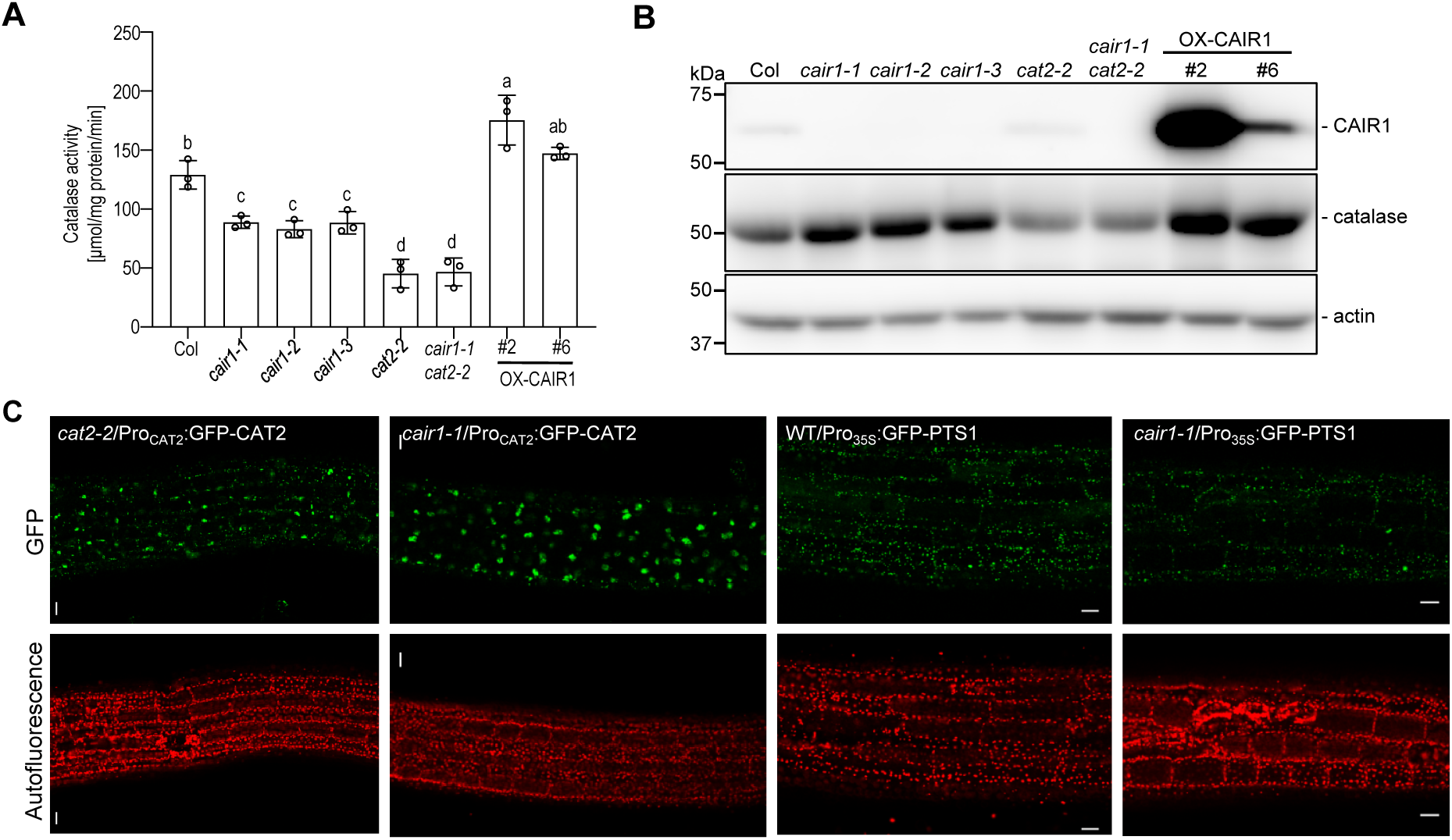
*cair1* show low catalase activity, associated with catalase aggregation. **(A)** Catalase activity in 5-d-old seedlings of wild type (Col), *cair1-1*, *cair1-2*, *cair1-3*, *cat2-2*, *cair1-1 cat2-2*, and two CAIR1 complementation lines (OX-CAIR1 #2 and #6). Data shown are means of 3 independent measurements. Shared letters indicate no statistically significant difference between the means (P > 0.05), as determined by two-way ANOVAs followed by Tukey’s test for multiple comparisons. **(B)** Immunoblot analysis of catalase protein levels in 5-d-old Col, *cair1-1*, *cair1-2*, *cair1-3*, *cat2-2*, *cair1-1 cat2-2*, and OX-CAIR1 #2 and #6 seedlings. Actin protein levels are shown as loading control. **(C)** GFP-CAT2 forms aggregates in peroxisomes when expressed in the *cair1-1* mutant background, but not in *cat2-2*. GFP-PTS1 serves as a control. The GFP signal was investigated in 7-d-old seedlings. Scale bars: 30 μm.

### *cair1* mutants show hypersensitivity to photorespiration stress and to alkaline growth conditions

Plant catalases have key roles in detoxifying H_2_O_2_ generated during photorespiration and fatty acid β-oxidation in peroxisomes (Baker *et al*, 2023). *cat2* mutants conditionally accumulate elevated H_2_O_2_ levels, which leads to activation of cell death in leaves, particularly when grown under long-day (LD) conditions (Kaurilind *et al*, 2015; Queval *et al*, 2007). In contrast to *cat2*, neither *cair1* mutants nor two *CAIR1*-overexpression lines (*cair1-1*/Pro_35S_:CAIR1; OX-CAIR1) displayed noticeable leaf lesions when grown in LD (Supplemental Figure S8A), a *cat2* mutant phenotype under LD associated with oxidative stress-induced salicylic acid (SA) signalling (Chaouch *et al*, 2010). Although macroscopic leaf lesions could not be detected in *cair1*, *cair1* and *cair1 cat2* showed elevated expression of the SA marker genes *PR1* and *PR2* under LD in comparison to wild type and *cat2*, respectively (Supplemental Figure S8B). Interestingly, however, under LD conditions with acute photorespiration stress, such as when 2-week-old plants grown in LD under high CO_2_ that inhibited photorespiration were transferred to air with ambient CO_2_ levels, *cair1* enhanced the *cat2* phenotype (Supplemental Figure S8C–E). Indeed, *cair1 cat2* displayed leaf lesions after 3 d following transfer to ambient CO_2_, whereas *cat2* leaf lesions were only detectable after 7 d (Supplemental Figure S8D, E). It is also noteworthy that the *cair1 cat2* double mutant displayed a smaller rosette size than *cat2* (Supplemental Figure S8D, E), suggesting reduced CAT1 and CAT3 activity in absence of CAIR1.

We further assessed if CAIR1 has a functional role under enhanced photorespiration and associated oxidative stress. We measured the maximum quantum yield of photosystem II (*F*v/*F*m) before and after 1 d of gas limitation (resulting in low CO_2_/high O_2_ conditions) causing enhanced photorespiration. Although less pronounced than *cat2*, *cair1* mutants displayed reduced *F*v/*F*m relative to the wild type under 24-h gas exchange limitation conditions that enhanced photorespiration (Figure 4A), whereas two CAIR1-overexpression lines conversely showed slightly higher *F*v/*F*m than wild type (Figure 4B). Moreover, *cair1 cat2* showed further reduced *F*v/*F*m relative to *cat2* under photorespiration-promoting conditions (Supplemental Figure S9A, B). ROS staining using 3,3′-diaminobenzidine (DAB) further indicated that *cair1* and *cat2* accumulated higher ROS levels compared to that in the wild type, whereas the CAIR1-overexpression line exhibited reduced ROS levels (Supplemental Figure S9C). In agreement with enhanced oxidative stress, *cair1* showed enhanced expression of two H_2_O_2_-inducible marker genes, namely *UGT74E2* and *GRX480* (Tognetti *et al*, 2010), after 24 h of enhanced photorespiration (Figure 4C, D).

**Figure 4.**
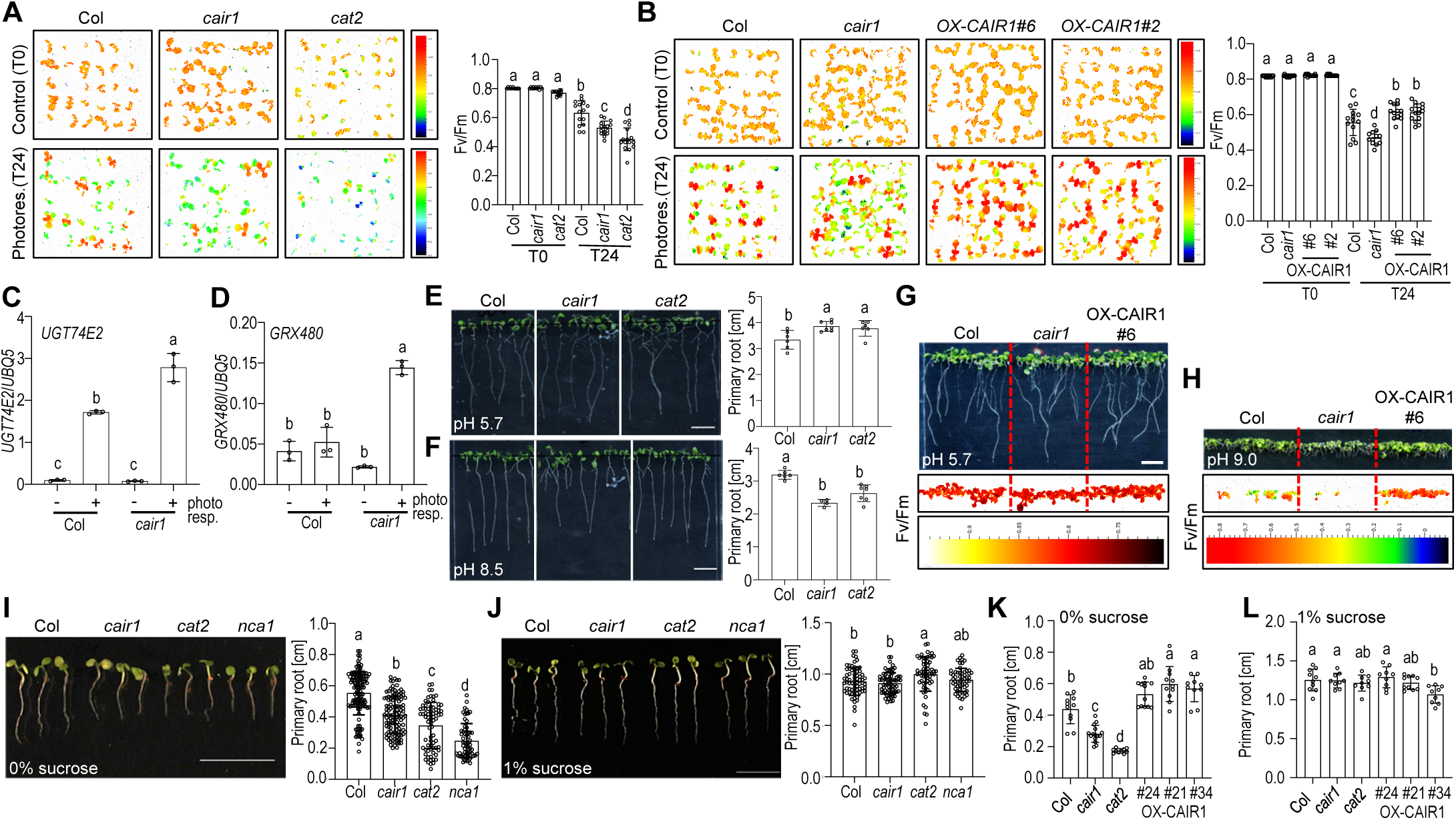
*cair1* show hypersensitivity to enhanced photorespiration and to high external pH. **(A, B)** Response to photorespiration–promoting growth conditions (gas limitation). (*Left*) False-color images representing *F*v/*F*m values of 10-d-old wild-type (Col), *cair1-1* (*cair1*), *cat2-2* (*cat2*), *cair1-1*/Pro_35S_:CAIR1 line #6 (OX-CAIR1 #6) and OX-CAIR1 #2 seedlings before [Control (T0)] and after 24 h of photorespiration–promoting growth conditions [Photores. (T24)]. *F*v/*F*m color scales are shown, with blue and red indicating low and high values, respectively. A representative plate is shown of triplicate replicate experiments. (*Right*) Quantitative *F*v/*F*m measurements. Independent plant data points and means ± SD (n >10) are shown. **(C, D)** RT-qPCR analysis of *UGT74E2* and *GRX480* gene expression in long day (LD)–grown, 7-d-old Col and *cair1* seedlings after 24 h of enhanced photorespiration (+) or control conditions (-). Independent experiment data points and means ± SD (n = 3) are shown. **(E, F)** Primary root length of seedlings at pH 5.7 (E) and pH 8.5 (F). Col, *cair1*, and *cat2* were grown for 5 d on plates containing ½ MS with 1% sucrose at pH 5.7, then transferred to ½ MS with 1% sucrose at either pH 5.7 or 8.5 and grown vertically. Photographs (*left*) and root length measurements (*right*) were taken 5 d and 8 d, respectively, after seedling transfer to differential pH. Independent plant data points and means ± SD (n > 5) are shown. Scale bars: 1 cm. **(G, H)** *cair1* is hypersensitive at pH 9.0. Five-d-old Col, *cair1*, and OX-CAIR1 #6 seedlings were transferred from ½ MS medium with 1% sucrose pH 5.8 to plates with pH 5.8 (G) or pH 9.0 (H). Photographs (*top*) and false-color images (*middle*) representing *F*v/*F*m values of 10-d-old seedlings 5 d after transfer to differential pH. Scale bars: 0.5 cm. **(I, J)** *cair1* and *cat2* show sucrose-dependent root elongation. Seedlings were grown for 5 d under LD conditions on ½ MS medium with (J) or without (I) 1% sucrose. Photographs of representative seedling growth (*left*) and root length measurements (*right*), with independent plant data points and means ± SD (n ≥ 60), are shown. **(K, L)** CAIR1 overexpression lines (OX-CAIR1 #24, #21, and #34) show longer primary roots than Col, *cair1*, and *cat2* when grown on sucrose-free medium. Independent plant data points and means ± SD (n > 20) are shown. (A–F, I–L) Shared letters indicate no statistically significant difference between the means (P > 0.05), as determined by two-way ANOVAs followed by Tukey’s test for multiple comparisons.

*nca1* mutants are more sensitive to alkaline growth conditions (high external pH) that induce the production of ROS in different cellular compartments, including peroxisomes (Li *et al*, 2015). To investigate this sensitivity in wild-type, *cair1*, and *cat2*, we transferred 4-d-old seedlings from pH 5.7 growth medium to pH 8.5 growth medium. Like *nca1-1* (Li *et al*, 2015), *cat2* and *cair1* showed reduced primary root growth compared to that in wild type at pH 8.5, which did not occur at pH 5.7 (Figure 4E, F). Moreover, at the higher pH of 9.0, survival of *cair1* was reduced in comparison to wild type (determined by absence of *F*v/*F*m measurements), whereas a CAIR1-overexpressing line showed enhanced alkalinity tolerance in the same conditions (Figure 4G, H). We conclude that, as a positive regulator of catalase activity, CAIR1 contributes to resilient growth under alkaline conditions.

During seedling establishment, fatty acid β-oxidation in peroxisomes generates H_2_O_2_ that must be efficiently removed by catalase activity. On sucrose-free growth medium where fatty acid β-oxidation must fuel growth, the loss of CAT2 function results in reduced root elongation (Liu *et al*, 2017; Yang *et al*, 2019). We thus compared the growth of wild type, *cat2*, *nca1, cair1*, and OX-CAIR1 seedlings on sucrose-free and sucrose-supplemented medium. After 5 d, *cat2*, *nca1*, and *cair1* seedlings exhibited significantly shorter primary roots than wild type specifically in the absence of sucrose (Figure 4I, J), whereas CAIR1-overexpressing lines showed longer primary roots under these conditions (Figure 4K, L). By contrast, exogenous sucrose fully restored root growth in all mutants (Figure 4J, L). These results further indicate that CAIR1 is required for catalase activity to tolerate H_2_O_2_ generated during fatty acid β-oxidation.

### CAIR1 is an unstable protein under oxidative stress

Chlorophyll fluorescence (*F*v/*F*m) measurements comparing oxidative stress responses of *cair1* and wild type suggested that CAIR1 played a functional role mainly during the first 24–48 h of enhanced photorespiration (Figure 5A). Interestingly, extended exposure to enhanced photorespiration as well as exogenous H_2_O_2_ treatment led to the degradation of endogenous CAIR1 (Figure 5B, C), which was blocked with proteasome inhibitors (Figure 5D). Moreover, we assessed the impact of additional stress factors on CAIR1 stability and observed that both salt and hydroxyurea (HU; a catalase inhibitor) (Juul *et al*, 2010) treatments destabilized CAIR1 (Figure 5E). By contrast, exogenous sucrose up to 2% (Liu *et al*, 2017) and CO_2_ supplementation, conditions known to reduce ROS production, both led to CAIR1 stabilization (Figure 5F, G). Taking these data together, we thus conclude that ROS destabilizes CAIR1 through proteasomal degradation.

**Figure 5.**
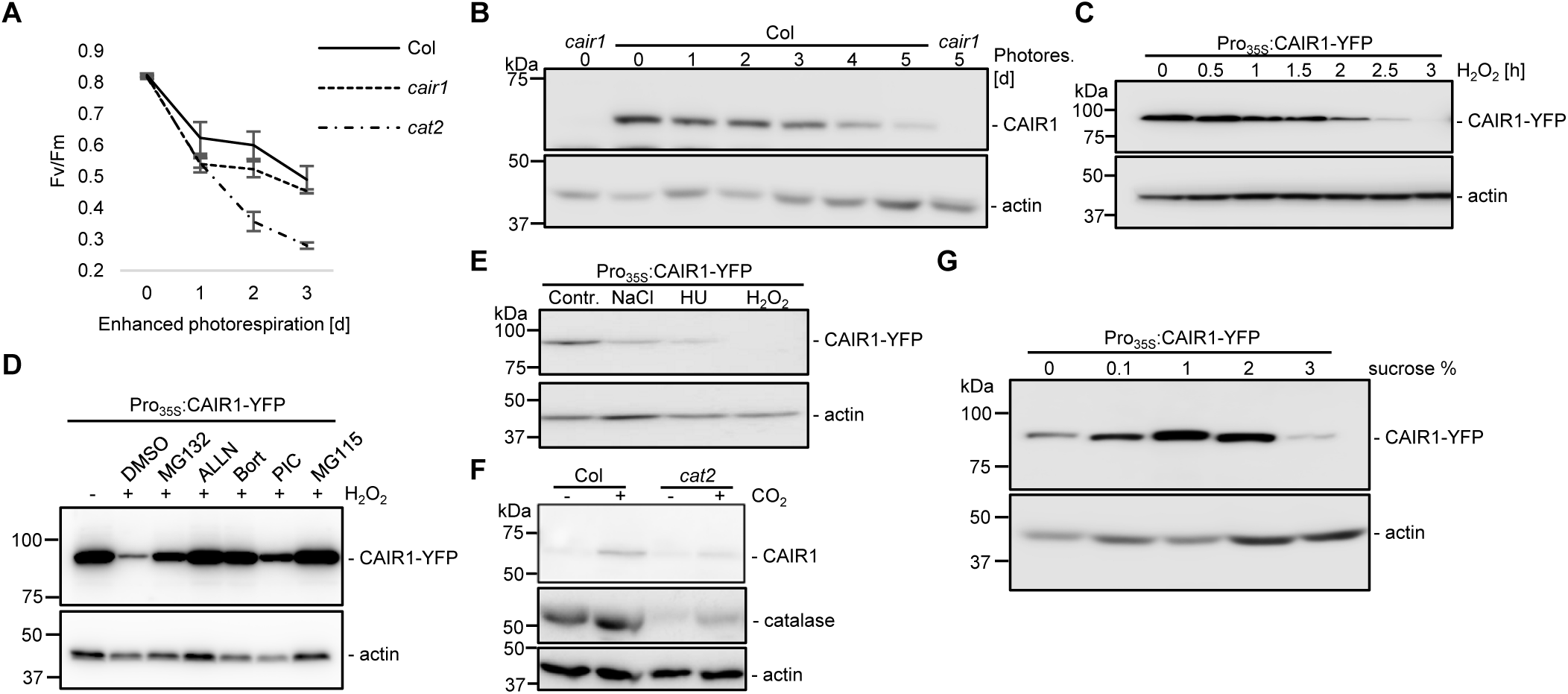
Proteasomal degradation of CAIR1 in response to ROS. **(A)** *cair1* mutant is sensitive to short-term enhanced photorespiration. *F*v/*F*m values of 10-d-old Col, *cair1-1* (*cair1*), and *cat2-2* (*cat2*) seedlings grown under long-day (LD) conditions following transfer to enhanced photorespiration conditions for the indicated time period [d]. **(B)** Endogenous CAIR1 proteins are degraded after 3 d of enhanced photorespiration treatment. Immunoblot analysis of endogenous CAIR1 in 6-d-old Col and *cair1* seedlings following their transfer from a CO₂-enriched chamber to conditions promoting enhanced photorespiration (Photores. [d]). **(C)** Exogenous H_2_O_2_ leads to destabilization of CAIR1-YFP protein. Immunoblot analysis of CAIR-YFP in 3-d-old *cair1-1*/Pro_35S_:CAIR1-YFP #23 (Pro_35S_:CAIR1-YFP) seedlings following their transfer to water supplemented with 20 mM H_2_O_2_ for the indicated time [h]. **(D)** Immunoblot analysis of CAIR-YFP in 3-d-old Pro_35S_:CAIR1-YFP seedlings that were pre-treated with 200 µM MG132, ALLN, bortezomib (Bort), protease inhibitor cocktail tablets (PIC), MG115, or DMSO (mock control) for 4 h in water followed by treatment with 20 mM H_2_O_2_ for 3 h (+ H_2_O_2_). **(E)** Salt stress, hydroxyurea, and H_2_O_2_ treatments similarly destabilize CAIR1. Immunoblot analysis of CAIR-YFP in 3-d-old Pro_35S_:CAIR1-YFP seedlings following their transfer to liquid MS medium supplemented with 100 mM NaCl (NaCl), 3 mM HU (HU), or 20 mM H_2_O_2_ (H_2_O_2_) compared to straight liquid MS (Contr.) for 3 h. **(F)** CO_2_ supplementation stabilizes CAIR1 levels. Immunoblot analysis of CAIR1 and catalase in 5-d-old Col and *cat2* seedlings grown in LD with (+) or without (-) CO_2_ supplementation. **(G)** The presence of sucrose stabilizes CAIR1-YFP protein level and inhibits oligomerization of CAIR1-YFP. Immunoblot analysis of CAIR-YFP in 6-d-old Pro_35S_:CAIR1-YFP seedlings cultured in media containing various sucrose concentrations (%). (B–G) Actin was used as loading control.

### Redox-dependent oligomerization of CAIR1 modulates its interaction with catalases

In Y2H and BIFC assays, CAIR1 showed interaction with itself, suggesting that CAIR1 oligomerizes *in vivo* (Figure 6A, B). Indeed, HA-CAIR1 co-immunoprecipitated with CAIR1-YFP in extracts from a Pro_35S_:CAIR1-YFP Pro_35S_:HA-CAIR1 double transgenic line (Figure 6C). Interestingly, anti-CAIR1 immunoblot analysis of extracts prepared under non-reducing conditions in the absence of dithiothreitol (DTT) detected bands at sizes approximate to that of CAIR1 dimers and higher order oligomeric forms that were absent in *cair1* (Figure 6D). Based on this observation, we further investigated whether CAIR1 oligomerization is dependent on its redox status by treating protein extracts from 5-d-old seedlings with varying concentrations of DTT as a reducing agent and H_2_O_2_ as an oxidizing agent. Notably, only the monomeric form of CAIR1 was detected in the presence of 10 and 20 mM DTT (Figure 6E). However, as oxidative conditions intensified, higher order oligomeric forms of CAIR1 emerged (at sizes approximate to that of homodimeric and -tetrameric forms), indicating that CAIR1 oligomerization is redox-dependent (Figure 6E). This response was specific for CAIR1, since, in sharp contrast to YFP-CAIR1, no redox-dependent oligomerization could be observed for YFP-HA, HPR1, GFP-CAT2, NCA1-GFP, or the related RCC1-like family members GFP-UVR8 and YFP-AT5G16040 (Supplemental Figure S10A–C). Consistent with its elevated endogenous ROS levels, *cat2* displayed enhanced CAIR1 oligomerization compared to that in wild type (Figure 6F). Similarly, enhanced photorespiration, which is associated with elevated levels of endogenous ROS, also increased CAIR1 oligomerization (Figure 6G).

**Figure 6.**
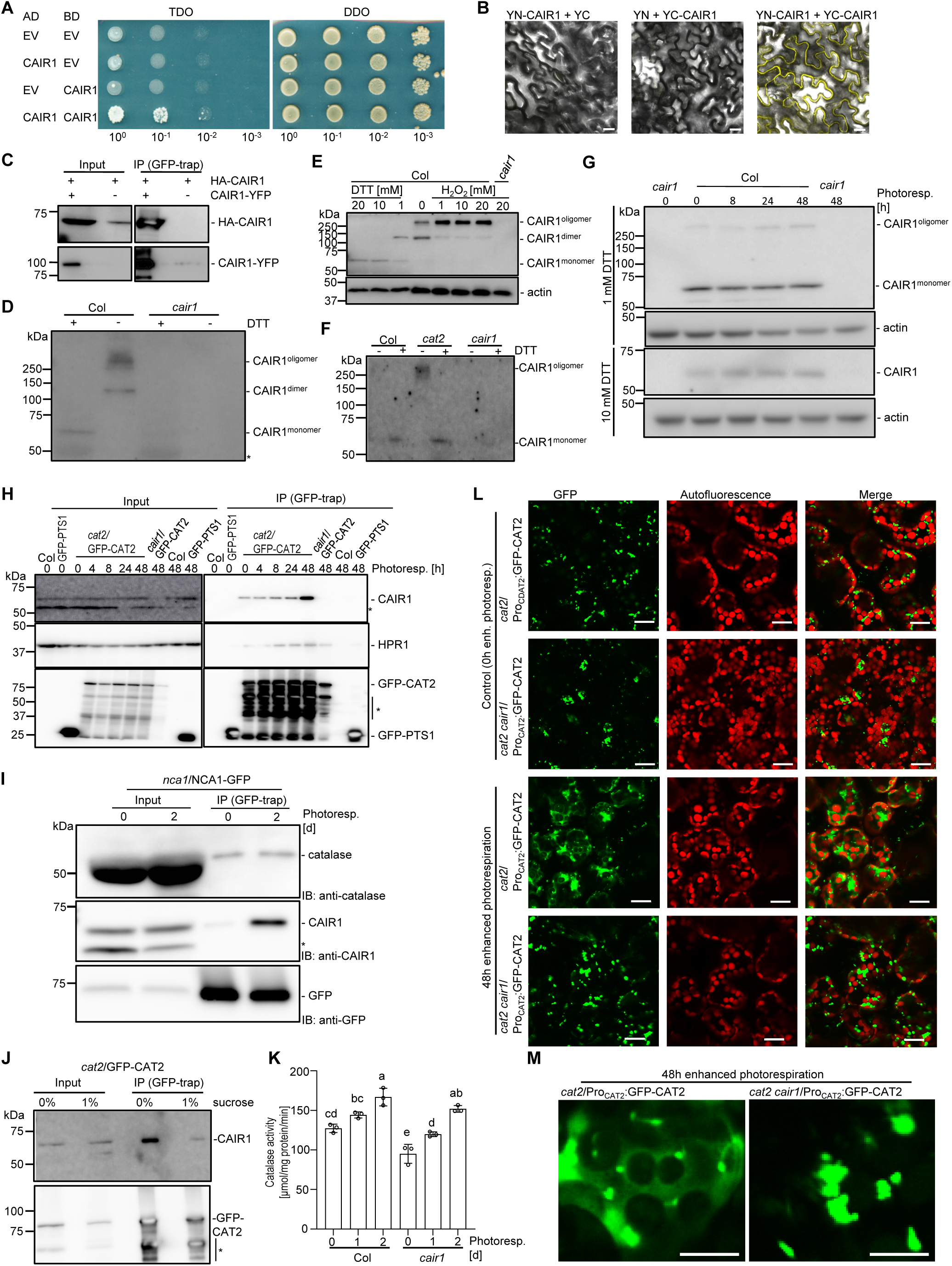
ROS facilitates the oligomerization of CAIR1 and regulates interaction with catalase. **(A)** Yeast two-hybrid analysis of CAIR1 and CAIR1 interactions with growth assays. AD, activation domain; BD, DNA binding domain; +His, +His medium (SD/Trp/Leu) as transformation control; -His, selective -His medium (SD/Trp/Leu/His); EV, empty vector. **(B)** BIFC assay in *N. benthamiana* leaves transiently expressing YFP fusion proteins YFPN-CAIR1 (YN-CAIR1) and YFPC-CAIR1 (YC-CAIR1) showing that YN-CAIR1 interacts with YC-CAIR1 in the cytosol. Scale bars: 30 μm. **(C)** Co-IP assay of CAIR1 fusion proteins using GFP-trap purification, showing that Pro_35S_:CAIR1-YFP interacts with Pro_35S_:3xHA-CAIR1 *in planta*. **(D)** Oligomerization of CAIR1 occurs without DTT. Immunoblot analysis of CAIR1 in 5-d-old wild-type (Col) and *cair1-1* (*cair1*) mutant seedlings with protein extraction performed with (+) or without (-) 5 mM DTT. **(E)** Redox level mediates the oligomerization of CAIR1. Immunoblot analysis of CAIR1 in 5-d-old Col and *cair1* seedlings with protein extractions initially performed without DTT, then treated with varying concentrations of DTT or H_2_O_2_ as indicated. Actin was used as a loading control. **(F)** *cat2* displays enhanced CAIR1 oligomerization compared to wild type under long-day (LD) conditions. Immunoblot analysis of CAIR1 in 5-d-old Col, *cat2-2* (*cat2*), and *cair1* seedlings grown in LD, with protein extractions performed with (+) or without (-) DTT. **(G)** Enhanced photorespiration increases CAIR1 oligomerization. Immunoblot analysis of CAIR1 in Col and *cair1* seedlings which were exposed to enhanced photorespiration conditions for different times (Photoresp. [h]), with proteins extracted with lysis buffer containing 1 mM or 10 mM DTT. Actin was used as a loading control. **(H)** ROS enhances interaction of CAIR1 and CAT2 *in planta*. Co-IP assay of CAIR1 and HPR1 proteins using GFP-trap purification. Alongside various control genotypes, 10-d-old *cat2-2*/Pro_CAT2_:GFP-CAT2 #6 (*cat2*/GFP-CAT2) seedlings grown under ambient CO_2_ conditions then subject to enhanced photorespiration conditions for various durations, with proteins extracted using lysis buffer lacking DTT. Asterisks (*) indicate likely degradation products. **(I)** Co-IP assay of catalase and CAIR1 proteins using GFP-trap purification, showing that ROS enhances interaction of NCA1 and CAIR1, but not interaction of NCA1and catalases *in planta*. Proteins were extracted from 10-d-old *nca1-1*/Pro_NCA1_:NCA1-GFP #11 (*nca1*/NCA1-GFP) seedlings before (0) and following 2 d enhanced photorespiration treatment (2) using lysis buffer without DTT. **(J)** Co-IP assay of CAIR1 using GFP-trap purification showing that exogenous sucrose inhibits the interaction of CAIR1 and catalase *in planta*. Proteins were extracted from 5-d-old *cat2-2*/GFP-CAT2 #2 (*cat2*/GFP-CAT2) seedlings grown on medium containing either 0% or 1% sucrose medium, using lysis buffer lacking DTT. Asterisks (*) indicate likely degradation products. **(K)** CAIR1 mediates ROS-enhanced catalase activity *in planta*. Proteins were extracted from 5-d-old Col and *cair1* seedlings grown under ambient CO_2_ conditions then subject to 0, 1, or 2 d of enhanced photorespiration conditions. Independent triplicate data points and means ± SD are shown. Shared letters indicate no statistically significant difference between the means (P > 0.05), as determined by two-way ANOVAs followed by Tukey’s test for multiple comparisons. **(L)** ROS changes catalase subcellular localization through CAIR1. *cat2-2*/Pro_CAT2_:GFP-CAT2 (*cat2*/Pro_CAT2_:GFP-CAT2) and *cat2-2 cair1-1*/Pro_CAT2_:GFP-CAT2 (*cat2 cair1*/Pro_CAT2_:GFP-CAT2) seedlings were grown for 10 d before exposure or not to enhanced photorespiration conditions for 2 d, after which GFP-CAT2 localization was investigated with a confocal microscope. Scale bar: 20 µm. **(M)** Higher magnification images of GFP-CAT2 localization in *cat2*/Pro_CAT2_:GFP-CAT2 and *cat2 cair1*/Pro_CAT2_:GFP-CAT2 after 2 d enhanced photorespiration conditions. Scale bars: 10 µm.

We further investigated whether redox-dependent oligomerization of CAIR1 influences its interaction with CAT2. Indeed, enhanced photorespiration increased interaction of CAIR1 with GFP-CAT2 and NCA1-GFP as determined by co-IP assays (Figure 6H, I), but did not affect the interaction between NCA1-GFP and catalases (Figure 6I). Interestingly, interaction of GFP-CAT2 with HPR1 also increased under these conditions (Figure 6H). We also found that exogenous sucrose, which reduces ROS production during seedling establishment (Liu *et al*, 2017), significantly inhibited interaction of GFP-CAT2 with CAIR1 (Figure 6J). We then assessed whether catalase activity is altered under growth conditions associated with elevated ROS levels. Catalase activity indeed increased under enhanced photorespiration in both wild type and *cair1*, although overall catalase activity remained lower in *cair1* (Figure 6K). Moreover, incubation under enhanced photorespiration conditions for 2 d resulted in the CAIR1-dependent, altered subcellular localization of GFP-CAT2, manifesting as enhanced cytosolic localization (Figure 6L, M). Our findings suggest that ROS reversibly regulates CAIR1 oligomerization state and its interaction with catalases, which affects catalase subcellular localization.

### Cys-356 and Cys-545 contribute to redox-regulated CAIR1 oligomerization and function

ROS-induced redox changes can oxidize cysteine thiols (RSH/RS-), producing highly reactive sulfenic acids (RSOH). These intermediates can then react with additional thiol groups to form disulfide bonds, promoting redox-dependent homo- or hetero-oligomerization of proteins (Cremers & Jakob, 2013; Huang *et al*, 2011; Iwona A. Buskiewicz, 2016; Roy *et al*, 2025). CAIR1 contains 12 cysteines (Supplemental Figure S11A), and mutation of each of these to alanine (CAIR1^12C-A^) abolished redox-dependent oligomerization in *N. benthamiana* transient assays and reduced it in Arabidopsis, thus supporting CAIR1 oligomerization via disulfide bond-formation (Supplemental Figure S11B and C). Among the 12 cysteines, AlphaFold3 structural analysis predicted 5 cysteines to be in the hydrophobic core and thus unlikely to be involved in redox regulation (C204, C270, C291, C330, C390), and 7 cysteines to be surface exposed (C36, C96, C116, C145, C356, C424, C545), with in particular Cys-356 and Cys-545 residues apparently well-conserved across plants (Supplemental Figure S11D). Interestingly, mutating them both to alanine (CAIR1^C356A,C545A^) reduced CAIR1 oligomerization in response to elevated ROS, as demonstrated in Arabidopsis and in *N. benthamiana* transient expression assays (Figure 7A, Supplemental Figure S11E). However, CAIR1^12C-A^ still showed ROS-mediated CAIR1 degradation and sucrose-mediated CAIR1 stabilization (Supplemental Figure S11F, G). We then tested whether cysteine-dependent oligomerization is required for the cellular redox-enhanced interaction between CAIR1 and catalase in Arabidopsis. In comparison to CAIR1, CAIR1^C356A,C545A^ and CAIR1^12C-A^ indeed exhibited reduced interaction with catalases under stress conditions (Figure 7B). Furthermore, expressing CAIR1^12C-A^ or CAIR1^C356A,C545A^ in *cair1-1* only partially rescued the *cair1* mutant phenotype under enhanced photorespiration conditions (Figure 7C). Together, these findings indicate that cellular redox status may regulate CAIR1 oligomerization involving Cys-356 and Cys-545, thereby modulating the CAIR1–catalase interaction and maintaining ROS homeostasis. However, ROS-mediated CAIR1 degradation occurs independently of its cysteine residues.

**Figure 7.**
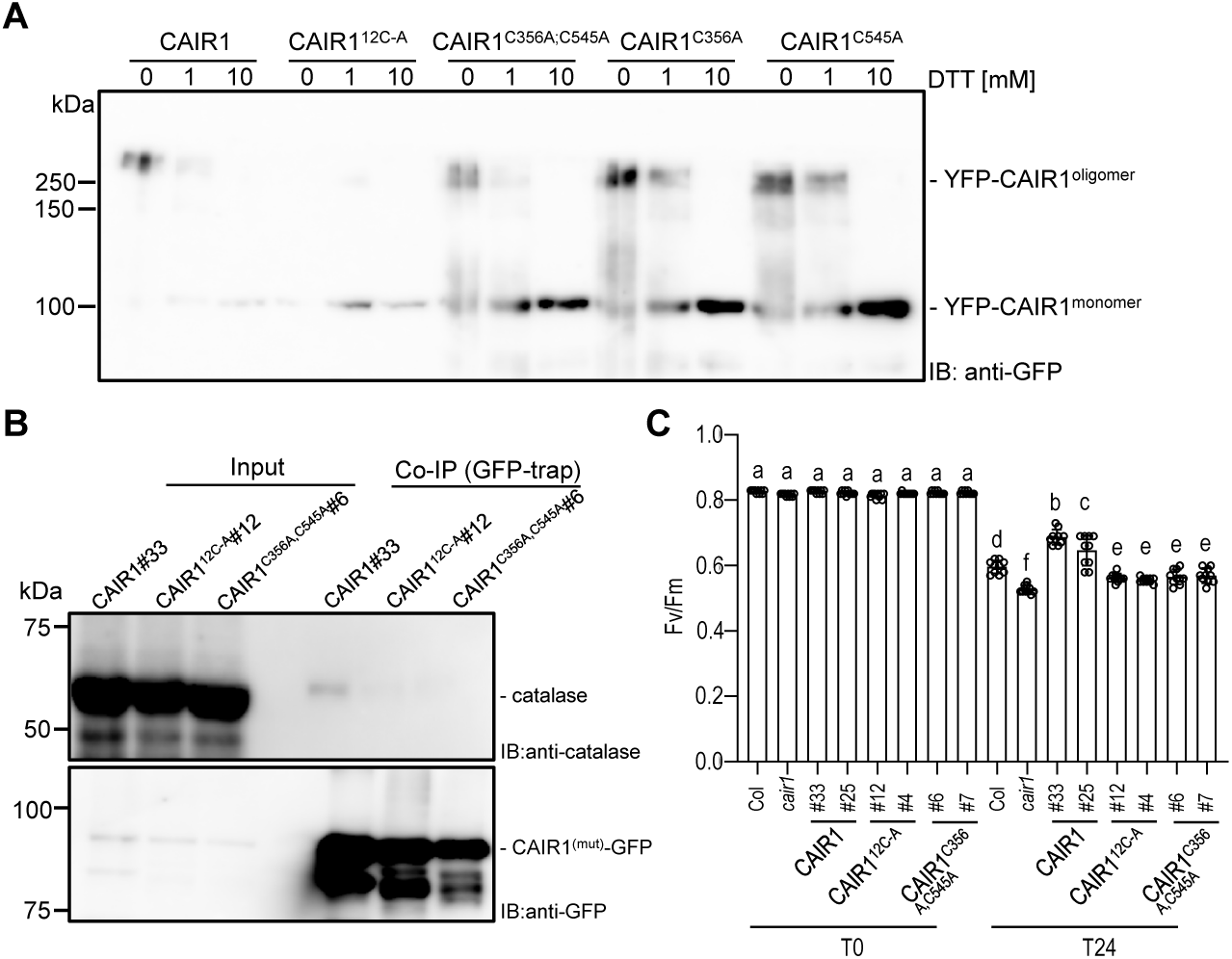
Cys-356 and Cys-545 contribute to CAIR1 function. **(A)** Immunoblot analysis of YFP-CAIR1, showing CAIR1 oligomeric state in 5-d-old seedlings expressing the CAIR1 variants CAIR1^12C-A^, CAIR1^C356A;C545A^, CAIR1^C356A^, or CAIR1^C545A^ with a YFP-tag under control of the 35S-promoter in the *cair1-1* background, with protein extractions performed with different concentrations of DTT. **(B)** Co-IP assay of catalase using GFP-trap purification, showing that cysteine mutation at C356 and C545 disturbs ROS-dependent interaction of CAT2 and CAIR1. Proteins were extracted from 10-d-old *cair1-1*/Pro_35S_:YFP-CAIR1 #33 (CAIR1#33), *cair1-1*/Pro_35S_:YFP-CAIR1^C12-A^ #12 (CAIR1^C12-A^#12), and *cair1-1*/Pro_35S_:YFP-CAIR1^C356A,C545A^ #6 (CAIR1^C356A,C545A^#6) seedlings exposed to 2 d enhanced photorespiration conditions using lysis buffer lacking DTT. **(C)** The cysteine of CAIR1 at C356 and C545 are required for oxidative response. Response to photorespiration-promoting growth conditions. *F*v/*F*m values of 10-d-old wild-type (Col), *cair1-1* (*cair1*), *cair1-1*/Pro_35S_:YFP-CAIR1 lines #25 and #33 (CAIR1 #25 and #33), *cair1-1*/Pro_35S_:YFP-CAIR1^C12-A^ lines #4 and #12 (CAIR1^C12-A^ #12 and #4), and *cair1-1*/Pro_35S_:YFP-CAIR1^C356A,C545A^ lines #6 and #7 (CAIR1^C356A,C545A^ #6 and #7) seedlings before (T0) and after 24 h of photorespiration–promoting growth conditions (T24). Independent data points and means ± SD are shown (n = 10). Shared letters indicate no statistically significant difference between the means (P > 0.05), as determined by two-way ANOVAs followed by Tukey’s test for multiple comparisons.

### CAIR1 mediates UV-B–induced inhibition of catalase activity

CAIR1, although identified as a previously undescribed interactor of the UVR8 photoreceptor, does not appear to influence UVR8-mediated UV-B-induced photomorphogenesis based on the standard phenotypic assays analyzed in this study. We therefore investigated whether UV-B alters CAIR1-mediated regulation of catalase activity. Wild-type plants grown in the presence of UV-B showed reduced catalase activity compared to those grown in absence of UV-B, whereas UV-B had no or a reduced effect on catalase activity in *cair1* and *uvr8* mutants, respectively (Supplemental Figure S12A). The UV-B effect on catalase activity was independent of any detectable effect on catalase protein abundance (Supplemental Figure S12B). However, plants grown under UV-B showed reduced interaction between CAIR1 and CAT2 in co-IP assays compared to plants grown in absence of UV-B (Supplemental Figure S12C). Interestingly, both endogenous CAIR1 and OX-CAIR1 protein levels increased under UV-B treatment (Supplemental Figure S12D,E). Collectively, these findings suggest that CAIR1 and UVR8 mediate UV-B–induced inhibition of catalase activity through suppression of the CAIR1–CAT2 interaction.

## Discussion

The interplay between ROS metabolism, peroxisomal function, and catalase activity is central to cellular redox homeostasis and stress responses. Catalases decompose H_2_O_2_ into water and molecular oxygen and their function in peroxisomes controls cellular H_2_O_2_ levels. In this study, we identified CAIR1 through its co-purification with GFP-UVR8 by AP–MS. CAIR1 is a previously functionally uncharacterized Arabidopsis RCC1-Like protein family member that interacts with catalases and NCA1 in the cytosol and positively regulates catalase activity, likely by regulating catalase entry into peroxisomes. CAIR1 undergoes reversible redox-dependent oxidation involving Cys356 and Cys545, resulting in changes in its oligomeric state. This redox-controlled oligomerization modulates CAIR1–catalase interactions and thereby fine-tunes catalase activity in response to the cellular redox environment (Figure 8).

**Figure 8.**
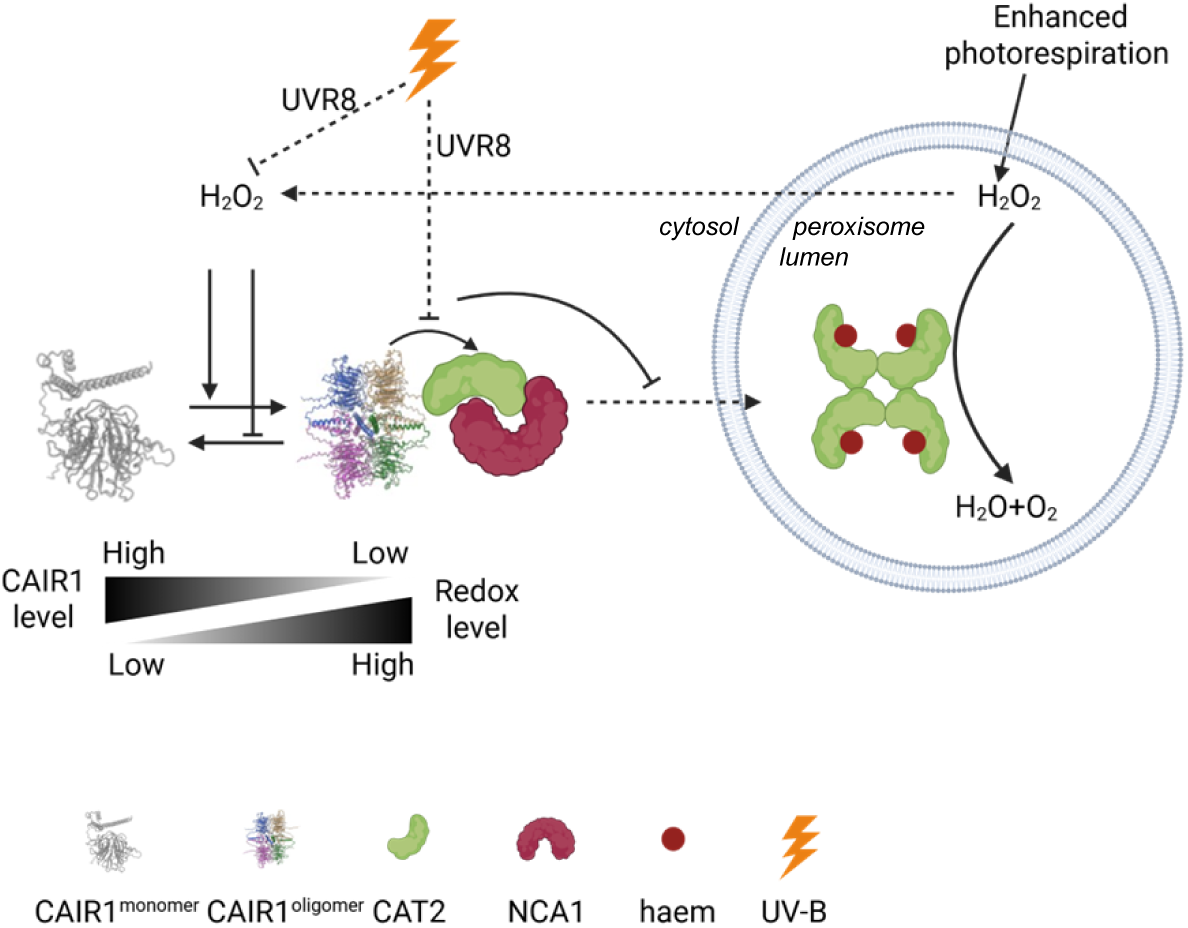
Working model of CAIR1-mediated regulation of catalase activity in Arabidopsis. Cellular ROS levels in Arabidopsis are tightly regulated through CAIR1 reversible oligomerization, which modulates catalase activity: (*Left of diagram*) Typically, catalase activity in peroxisomes converts potentially damaging H_2_O_2_ to benign H_2_O and O_2_. Elevated H_2_O_2_ in peroxisomes results from enhanced photorespiration. The mechanism by which H_2_O_2_ is transmitted between compartments, such as from peroxisomes to the cytosol or other organelles, remains unknown (indicated as dashed arrow). Elevated H_2_O_2_ in the cytosol promotes CAIR1 oligomerization (indicated as CAIR1^oligomer^), as mediated by the C356 and C545 residues, and accelerates CAIR1 degradation. CAIR1 oligomerization enhances CAIR1 interaction with catalases (CAT2) and its chaperone NCA1 in the cytoplasm, thereby inhibiting catalase peroxisomal import and increasing catalase activity in the cytosol. (*Top right of diagram*) Under UV-B, UVR8 interaction with CAIR1 mediates inhibition of catalase activity by weakening the interaction between CAIR1 and CAT2. It remains unclear whether UV-B regulates ROS levels through UVR8 to indirectly suppress the CAIR1–CAT2 interaction, or whether UV-B directly inhibits the CAIR1–CAT2 interaction in a UVR8-dependent manner (as indicated by dashed inhibitory lines). Overall, these molecular interactions result in an inverse relationship between CAIR1 level and redox level in the cytosol, which is inversely regulated by UV-B-activated UVR8. Created in BioRender (https://app.biorender.com/).

Although the importance of catalases in peroxisomes is well established, a mechanistic understanding of their peroxisomal import remains incomplete. Plant catalases lack a canonical C-terminal PTS1 motif, yet their import depends on PEX5, the receptor for PTS1-containing matrix proteins (Kamigaki *et al*, 2003; Mhamdi *et al*, 2012; Oshima *et al*, 2008), which is consistent with our observation that PEX5 co-purified with GFP-CAT2 (Supplemental Table S1). Studies in fungi have shown that weak, non-canonical PTS1 signals delay peroxisomal import, thus allowing proper folding and preventing aggregation of catalase within the peroxisomal matrix (Williams *et al*, 2012). The same may apply in plants, given that artificial addition of a canonical PTS1 to Arabidopsis CAT2 failed to fully restore catalase activity or mutant phenotypes (Al-Hajaya *et al*, 2022). Although plant non-canonical PTS1 signals differ from those in fungi and mammals and remain poorly defined (Mhamdi *et al*, 2012), the available evidence indicates that import kinetics are critical for catalase functionality.

Consistent with the model that catalase import kinetics are important, we observed enhanced aggregation of GFP-CAT2 in *cair1* mutants, whereas the localization of GFP-PTS1 was unaffected. This suggests that CAIR1 forms a cytosolic complex with CAT2 that regulates the rate of catalase import into peroxisomes, which appears essential for proper enzymatic activity. Notably, PEX5 was detected only in GFP-CAT2 AP–MS, but not in CAIR1-GFP or NCA1-GFP datasets, suggesting that CAIR1 and NCA1 transiently associate with CAT2 in the cytosol and thus delay its interaction with PEX5, preventing premature peroxisomal import. Thus, basal catalase activity likely depends on tightly controlled import kinetics mediated by CAIR1.

Under conditions leading to enhanced photorespiration, cytosolic accumulation of CAT2 was strictly CAIR1 dependent (Figure 6L,M). This may reflect direct CAIR1-mediated retention of CAT2 in the cytosol under elevated ROS or indirect modulation of the peroxisomal import machinery. In support of redox-sensitive import control, PEX5 activity has been shown to decrease under oxidative stress in yeast (Apanasets *et al*, 2014; Walton *et al*, 2017). Similarly, in mammalian cells, oxidative stress induces phosphorylation of Pex14, a core component of the peroxisomal translocation machinery, thus impairing Pex5–Pex14 interactions, suppressing catalase import, and increasing cytosolic catalase levels (Dubreuil *et al*, 2020; He *et al*, 2021; Okumoto *et al*, 2020). These observations suggest that regulated cytosolic localization of catalases is evolutionarily conserved, although the molecular mechanisms differ among organisms. How CAIR1 mechanistically mediates catalase retention in the cytosol, and the relative contributions of cytosolic versus peroxisomal catalase to oxidative stress tolerance in plants, remain open questions.

In addition to redox-dependent regulation, our data reveal that CAIR1 also mediates UV-B–induced inhibition of catalase activity. Although CAIR1 interacts with the UV-B photoreceptor UVR8, it does not appear to participate in canonical UVR8-dependent photomorphogenic responses. Instead, UV-B exposure suppresses catalase activity in a manner that requires both CAIR1 and UVR8, without substantially affecting catalase protein abundance. This reduction in activity is accompanied by a UV-B-induced decrease in the CAIR1–CAT2 interaction, suggesting that disruption of this complex is a key mechanism underlying catalase inhibition. Notably, UV-B treatment stabilizes CAIR1 protein levels in a UVR8-dependent manner; however, the biological relevance of this stabilization remains unclear. Together, these findings identify a UV-B–responsive regulatory module in which UVR8 and CAIR1 collectively attenuate catalase activity. An important unresolved question is whether UV-B-activated UVR8 modulates cellular ROS homeostasis, such as via activation of antioxidant pathways, and indirectly weakens the CAIR1–CAT2 interaction or whether UVR8 directly interferes with the CAIR1–CAT2 complex upon UV-B perception.

Beyond their cytotoxic effects, ROS function as key signalling molecules in plant development and stress responses (Mhamdi & Van Breusegem, 2018; Mittler *et al*, 2022). Fluctuations in cellular redox status in response to environmental stresses affect protein functions through redox-dependent oxidation (Chi *et al*, 2013; Garcia-Santamarina *et al*, 2014; Roy *et al*, 2025; Tian *et al*, 2018; Wei *et al*, 2024). For example, activity and stability of redox-sensitive transcription factors are affected by the redox state of the cell (He *et al*, 2018; Lee *et al*, 2021; Shaikhali *et al*, 2008; Shaikhali *et al*, 2012; Tian *et al*, 2018; Wei *et al*, 2024). Given its central role in H_2_O_2_ homeostasis, catalase localization and activity must be tightly regulated (Foyer *et al*, 2020). Our data indicate that CAIR1, together with the catalase chaperone NCA1, assembles into a cytosolic regulatory complex that fine-tunes catalase activity. Our findings suggest that CAIR1 regulation operates through two distinct redox-sensitive layers. First, under rising cellular H_2_O_2_ levels, CAIR1 responds to cellular redox changes possibly through intermolecular disulfide bond formation involving Cys356- and Cys545, driving the formation of CAIR1 oligomers, thereby enhancing its interaction with catalases and subsequent peroxisomal import rates. Second, during prolonged oxidative stress CAIR1 protein abundance decreases, suggesting that higher or chronic ROS levels might initiate post-translational destruction pathways, possibly involving oxidation-dependent ubiquitination. As a result, CAIR1 levels drop when ROS levels are excessive or sustained over time, limiting its ability to facilitate catalase activation. Together, this dual mechanism enables plants to dynamically adjust their H_2_O_2_ scavenging capacity and maintain redox homeostasis under fluctuating environmental conditions.

AP–MS–based interactome analysis further expands our understanding of the protein networks associated with catalases. Our GFP-CAT2 AP–MS confirmed known interactors and identified numerous further candidates, including many peroxisomal enzymes. This is consistent with early ultrastructural observations of enzyme-rich crystalline cores within peroxisomes (Lodhi & Semenkovich, 2014; Rhodin, 1958). Notably, loss of CAT2 affects the activities of multiple peroxisomal enzymes in addition to catalase, including ACX, GOX, enoyl-CoA hydratase (ECH), L-hydroxyacyl-CoA dehydrogenase (HAD), KAT, isocitrate lyase (ICL), and malate synthase (MLS) (Eastmond, 2007; Liu *et al*, 2017; Yang *et al*, 2025; Yuan *et al*, 2017), although whether these effects are direct or indirect remains unclear.

In conclusion, CAIR1 is a previously undescribed cytosolic interactor of catalases that, likely in coordination with catalase maturation by NCA1, modulates catalase import dynamics into peroxisomes, thereby positively regulating catalase activity. The interaction with catalases is fine-tuned by reversible oligomerization of CAIR1, a process apparently governed by the cellular redox state. These findings suggest that CAIR1 functions as a ROS-responsive regulator, contributing to H_2_O_2_ homeostasis by modulating catalase activity under varying environmental conditions.

## Materials and Methods

### Plant material and growth conditions

*cair1-1* (GABI_438B07), *cat2-2* (SALK_076998) (Queval *et al*, 2007), *nca1-1* (N68673) (Hackenberg *et al*, 2013), *uvr8-6* (SALK_033468) (Favory *et al*, 2009), and *rup1 rup2* (Gruber *et al*, 2010) mutant lines as well as the Pro_35S_:GFP-PTS1 (Mano *et al*, 2002), sfGFP1-10^OPT^ targeted to peroxisomes (PX-sfGFP1-10; Pro_UBQ10_:sfGFP1-10^OPT^-SKL), cytoplasm (CYTO-sfGFP1-10; Pro_UBQ10_:sfGFP1-10^OPT^), nuclei (NU-sfGFP1-10; Pro_UBQ10_:sfGFP1-10^OPT^-NLS), and plasma membrane (PM-sfGFP1-10; Pro_UBQ10_:PIP2:sfGFP1-10^OPT^) (Park *et al*, 2017), and Pro_35S_:StrepII-3xHA-YFP transgenic lines (Lapin *et al*, 2019) are all in the Columbia (Col) background. *cair1-1 cat2-2* was generated by genetic crossing and genotyped by PCR (Supplemental Table S2).

As the T-DNA in *cair1-1* is in an intron and a second isoform of *CAIR1* is predicted in its second gene model (AT5G11580.2), we additionally generated two *cair1* null mutant alleles using CRISPR-Cas9. The guide oligonucleotides (Supplemental Table S2) were cloned into the binary vector pHEE401E binary vector following the published method (Wang *et al*, 2015). The resulting constructs pHEE401E-CAIR1#1 and pHEE401E-CAIR1#2 were transformed into the wild type and *cair1-1* background, respectively. *cair1-2*/*at5g11580-2* contains a 278-bp deletion in the 5^th^ exon in the *cair1-1* mutant background (thus knocking out both AT5G11580.1 and potential AT5G11580.2), and *cair1-3/at5g11580-3* contains an early frameshift mutation due to a single base pair insertion at position 92 after the start ATG (Supplemental Figure S3).

For AP–MS experiments, plants were grown on soil under long-day (LD) conditions in a GroBank growth chamber (CLF Plant Climatics) (120 µmol m^-2^ s^-1^ white light, with or without 1.5 µmol m^-2^ s^-1^ UV-B) (Arongaus *et al*, 2018). After three weeks, the plants exposed to UV-B were subjected to an additional 15 min of broadband UV-B irradiation before rosette tissue was harvested.

For experiments under aseptic conditions *in vitro*, Arabidopsis seeds were surface sterilized with chlorine gas and sown on half-strength Murashige & Skoog basal salt (MS) medium containing 0.5% or 0.3% (w/v) phytagel (Sigma) without sucrose. After 2 d of stratification at 4°C, seedlings were grown under LD conditions in a standard growth chamber (MLR-350, Sanyo; 16-h/8-h light/dark cycle) with 66 µmol m^-2^ s^-1^ white light (measured with a LI-250 light meter; LI-COR Biosciences).

For catalase lesion, seeds were sown on soil and stratified for 2 d at 4°C before transfer to LD conditions (16-h/8-h light/dark cycle, 120 µmol m^-2^ s^-1^ white light) in a CLF GroBank growth chamber at ambient 420 ppm CO_2_ or in an Incrementum 1400 (Bronson Climate BV) supplemented with CO_2_ at 3,000 ppm.

To induce oxidative stress through enhanced photorespiration, Petri dishes with plants grown on half-strength MS medium without sucrose for 7 d were sealed with three layers of Parafilm M (Amcor) to restrict the ambient air influx and exposed to continuous light (200 µmol m^-2^ s^-1^ white light) to prevent the confounding effects of night-time respiration on gas homeostasis.

### Plasmid construction and generation of transgenic plants

The coding sequences of *CAIR1*, *CAT1*, *CAT2*, *CAT3*, *NCA1*, and *HPR1* were amplified with corresponding primers respectively (Supplemental Table S2) from cDNA and cloned into entry vector pDONR207 using Gateway technology (ThermoFisher Scientific) to produce pDONR207-CAIR1-no stop codon, pDONR207-CAIR1, pDONR207-CAT1, pDONR207-CAT2, pDONR207-CAT3, pDONR207-NCA1, and pDONR207-HPR1. Cysteine mutant versions CAIR1 were constructed using a two-step PCR-based site-directed mutagenesis protocol. First the 5′-segment of CAIR1 cDNA was amplified with the CAIR1GWF primer and site-directed mutagenesis primers, which introduced Cysteine (C) to Alanine (A) amino acid exchanges. The PCR amplified and gel purified 5′ cDNA segments were then used as 5′ primers with the CAIR1GWR primer to amplify the mutagenized full-length CAIR1 C-A cDNA versions, which were then cloned into pDONR207 with Gateway technology. This step was then repeated to produce pDONR207-CAIR1^12C-A^ and pDONR207-CAIR1^C356A;C545A^.. To construct AD-CAIR1, BD-CAT1, BD-CAT2, BD-CAT3, and BD-NCA1, the entry vectors pDONR207-CAIR1, pDONR207-CAT1, pDONR207-CAT2, pDONR207-CAT3, and pDONR207-NCA1 were cloned with Gateway technology into pGADT7-GW-rfb (Lu *et al*, 2010) and pGBKT7-GW (Lu *et al*, 2010), respectively. To construct YFPN-CAT2, YFPC-HPR1, and YFPC-CAIR1, pDONR207-CAIR1, pDONR207-CAT2, and pDONR207-HPR1 were cloned into YFN43 (Belda-Palazon *et al*, 2012) and YFC43 (Belda-Palazon *et al*, 2012) with Gateway technology. To construct cLUC-CAT2, cLUC-NCA1, and CAIR1-nLUC, the coding sequence of CAT2, NCA1, and CAIR1 was obtained using corresponding primers, then cloned into pCAMBIA1300-cLUC (Chen *et al*, 2008) and pCAMBIA 1300-nLUC (Chen *et al*, 2008) with Gibson cloning, respectively. To construct Pro_35S_:CAIR1-YFP, Pro_35S_:YFP-CAIR1, Pro_35S_:YFP-CAIR1^12C-A^, Pro_35S_:YFP-CAIR1^C356A;C545A^, and Pro_35S_:CAIR1, pDONR207-CAIR1-no stop codon, pDONR207-CAIR1, pDONR207-CAIR1^12C-A^, and pDONR207-CAIR1^C356A;C545A^ were cloned into Pro_35S_:R1-R2-EYFP-Tnos (pGWB541), Pro_35S_:EYFP-R1-R2-Tnos (pGWB542), Pro_35S_:R1-R2-Tnos (pGWB402), and Pro_35S_:3xHA-R1-R2-Tnos (pGWB415) (Nakagawa *et al*, 2007), respectively, with Gateway technology.

To generate Flag-CAT2, Flag-NCA1, GFP-CAIR1, and UVR8 constructs for expression in HEK293 cells, the coding sequences of *CAT2* and *NCA1* were cloned into the p3xFLAG-CMV-10 vector (Life Science Market). The coding sequence of *CAIR1* was cloned into the pEGFP-C1 vector (Axybio). The coding sequence of *UVR8*, including the stop codon, was cloned into the pEGFP-N1 vector (Axybio). All constructs were generated using Gibson assembly, and all primers used for cloning are listed in Supplemental Table S2.

To construct Pro_CAIR1_:CAIR1-GFP, three fragments were first amplified and cloned into entry vectors using MultiSite Gateway Recombination Cloning Technology (primers listed in Supplemental Table S2). A fragment containing the 5ʹ regulatory sequence of *CAIR1* and the *CAIR1* genomic sequence (2,846 bp) was amplified and cloned into pDONR P4-P1R, the coding sequence of GFP including stop codon was amplified from N-SpR-GFP plasmid (Hu *et al*, 2019) and cloned into pDONR221, and a fragment containing *CAIR1* 3ʹ regulatory sequence (2,973 bp) was amplified and cloned into pDONR P2R-P3.

To construct Pro_NCA1_:NCA1-GFP, two fragments were first amplified and cloned into Gateway entry vectors (primers listed in Supplemental Table S2). A fragment containing the 5ʹ regulatory sequence and genomic sequence of NCA1 (3,666 bp) was amplified and cloned into pDONR P4-P1R and a fragment containing NCA1 3ʹ regulatory sequence (1,587 bp) was amplified and cloned into pDONR P2R-P3.

To construct Pro_CAT2_:GFP-CAT2, three fragments were first amplified and cloned into Gateway entry vectors (primers listed in Supplemental Table S2). A fragment containing the 5ʹ regulatory sequence of *CAT2* (943bp) was amplified and cloned into pDONR P4-P1R, the coding sequence of GFP without stop codon was amplified from N-SpR-GFP plasmid (Hu *et al*, 2019) and cloned into pDONR221, and a fragment containing the *CAIR1* genomic sequence including 3ʹ regulatory sequence (3,203 bp) was amplified and cloned into pDONR P2R-P3.

The entry vectors pDONR P4-P1R, pDONR221, and pDONR P2R-P3 carrying the above relevant fragments to generate Pro_CAIR1_:CAIR1-GFP, Pro_NCA1_:NCA1-GFP, and Pro_CAT2_:GFP-CAT2 genomic clones, respectively, were cloned into pFR7m34GW (Kalmbach *et al*, 2023) by MultiSite Gateway cloning technology.

To construct Pro_UVR8_:GFP-UVR8, an *E. coli* strain with BAC clone MGI19 carrying the *UVR8* gene was obtained from the Arabidopsis Biological Resource Center (ABRC). BAC recombineering (Hu *et al*, 2019) was used to add a GFP tag at the N terminus of UVR8 within the *UVR8* genomic clone.

To construct CAIR1-sfGFP11, the coding sequence of *CAIR1* was amplified from cDNA with primers CAIR1F and CAIR1superfoldGFPR (Supplemental Table S2) and cloned into entry vector pDONR207 using Gateway technology (ThermoFisher Scientific) to produce pDONR207-CAIR1-sfGFP11. Then pDONR207-CAIR1-sfGFP11 was cloned into Pro_35S_:R1-R2-Tnos (pGWB402) (Nakagawa *et al*, 2007).

All resulting binary vectors were introduced into agrobacterial strain GV3101 and transformed into the corresponding mutant background or wild type by the floral dip method (Clough & Bent, 1998). Homozygous lines with single-locus insertion were identified and used in the T3 generation.

### Immunoblot analysis, co-immunoprecipitation

Rabbit polyclonal antibodies were generated against a synthetic peptide derived from the CAIR1 protein sequence (amino acids 373–387: C+RKISGGSSRFRDPVQ) and were affinity purified against the peptide (Eurogentec).

For immunoblot analysis, total protein extracts from seedlings were prepared using a previously described method (Hu *et al*, 2019). Briefly, samples were homogenized in extraction buffer (50 mM Tris-HCl [pH 7.5], 10% [v/v] glycerol, 1 mM EDTA, 150 mM NaCl, 0.1% [v/v] Triton X-100) supplemented with freshly added 5 mM DTT, 1 mM PMSF, and 20 µM Sigma plant protease inhibitor cocktail. Protein concentrations were determined using the Bradford assay, following the manufacturer’s instructions. Aliquots of 20 µg protein were separated on 10% SDS-polyacrylamide gels and transferred to PVDF membranes using the iBlot 2 Dry Blotting System. GFP and YFP fusion proteins were detected using the Living Colors® GFP Monoclonal Antibody (Clontech, dilution 1:5000) in TBST buffer. Endogenous catalase and HPR1 proteins were detected using anti-catalase (Agrisera, product no. AS09 501) and anti-HPR (Agrisera, product no. AS11 1797) antibodies in PBST and TBST buffers, respectively, following the manufacturer’s protocols. Horseradish peroxidase (HRP)–conjugated anti-rabbit and anti-mouse immunoglobulins (Dako A/S) were used as secondary antibodies. Protein bands were visualized using ECL Select Western Blotting Detection Reagent (GE Healthcare) and analyzed with an Amersham Imager 680 camera system (GE Healthcare). For loading control, the membrane was stripped with 0.2 M NaOH for 20 minutes and then probed with anti-actin antibody (Sigma).

For co-IP and affinity purification, protein extraction was performed as described above for immunoblot analysis. Total protein extracts were incubated with 40 µL GFP-Trap agarose (ChromoTek) affinity resin for 1 h at 4°C, after ensuring equal protein concentrations. The GFP-Trap was then washed four times with washing buffer (50 mM Tris-HCl [pH 7.5], 150 mM NaCl). For co-immunoprecipitation, 40 µL of 1x Laemmli buffer was added to the samples, which were then boiled at 98°C for 7 min. The samples were subsequently separated by SDS-PAGE for further analysis.

### Affinity purification–mass spectrometry (AP–MS)

Approximately 15 g of freshly harvested leaf tissue per sample were ground in liquid nitrogen and extracted with 30 mL of lysis buffer (50 mM Tris-HCl [pH 7.5], 10% [v/v] glycerol, 1 mM EDTA, 150 mM NaCl, 1% [v/v] Triton X-100). The extracts were centrifuged at full speed (7,200 *g*) for 20 min at 4°C. From each sample’s supernatant, technical triplicates of 20 mg of total protein each were generated and adjusted to an equal volume of 10 mL in lysis buffer and used for immunoprecipitation. The extracts were incubated for 1 h at 4°C with 50 µL GFP-Trap® Agarose resin (ChromoTek), which had been prewashed three times with 1.5 mL extraction buffer and resuspended in 1 mL lysis buffer before addition to the protein samples. The resins were pelleted at 500 rpm at 4°C and washed four times with 10 mL washing buffer (50 mM Tris–HCl [pH 7.5], 300 mM NaCl). Then, the resin was resuspended in 1 mL washing buffer, transferred to 1.5-mL Eppendorf tubes, and pelleted again. As controls, protein extracts were prepared in triplicate from wild type (Col) and Pro_35S_:StrepII-3xHA-YFP (Lapin *et al*, 2019) plants of the same age, harvested and processed in parallel with the Pro_UVR8_:GFP-UVR8, Pro_35S_:CAIR1-YFP, Pro_35S_:YFP-CAIR1, Pro_CAIR1_:CAIR1-GFP, and Pro_NCA1_:NCA1-GFP samples. For GFP-CAT2 purification, the co-purifying proteins obtained with the peroxisomal marker line Pro_35S_:GFP-PTS1 (Mano *et al*, 2002) were included as an additional control Sample preparation and LC-MS/MS data acquisition: Enriched proteins were submitted to an on-bead digestion using trypsin. In brief, dry beads were re-dissolved in 25 µL digestion buffer 1 (50 mM Tris, pH 7.5, 2M urea, 1mM DTT, 5 ng/µL trypsin) and incubated for 30 min at 32°C in a Thermomixer with 400 rpm. Next, beads were pelleted and the supernatant was transferred to a fresh tube. Digestion buffer 2 (50 mM Tris, pH 7.5, 2M urea, 5 mM CAA) was added to the beads, and after mixing the beads were pelleted and the collected supernatant was combined with the previous one. The combined supernatants were then incubated o/n at 32°C in a Thermomixer set to 400 rpm, with samples protected from light during incubation. The digestion was stopped by adding 1 µL TFA and desalted with C18 Empore disk membranes according to the StageTip protocol (Rappsilber *et al*, 2003).

Dried peptides were re-dissolved in 2% (v/v) ACN, 0.1% (v/v) TFA (10 µL) for analysis. Samples were analyzed using an EASY-nLC 1200 (Thermo Fisher) coupled to a Q Exactive Plus mass spectrometer (Thermo Fisher) or using an EASY-nLC 1000 (Thermo Fisher) coupled to a Q Exactive mass spectrometer (Thermo Fisher). Peptides were separated on 16-cm frit-less silica emitters (New Objective, 75-µm inner diameter), packed in-house with reversed-phase ReproSil-Pur C18 AQ 1.9 µm resin (Dr. Maisch, High Performance LC GmbH). Peptides were loaded on the column and eluted for 115 min using a segmented linear gradient of 5% to 95% solvent B (0 min: 5% B; 0–5 min: 5% B; 5–65 min: 20% B; 65–90 min: 35% B; 90–100 min: 55% B; 100–105 min: 95% B; 105–115 min: 95% B) (solvent A 0% ACN, 0.1% [v/v] FA; solvent B 80% [v/v] ACN, 0.1% [v/v] FA) at a flow rate of 300 nL/min. Mass spectra were acquired in data-dependent acquisition mode with a TOP15 method. MS spectra were acquired in the Orbitrap analyzer with a mass range of 300–1,750 m/z at a resolution of 70,000 FWHM and a target value of 3×10^6^ ions. Precursors were selected with an isolation window of 1.3 m/z (Q Exactive Plus) or 2.0 m/z (Q Exactive). HCD fragmentation was performed at a normalized collision energy of 25. MS/MS spectra were acquired with a target value of 105 ions at a resolution of 17,500 FWHM, a maximum injection time (max.) of 55 ms and a fixed first mass of m/z 100. Peptides with a charge of +1, greater than 6, or with unassigned charge state were excluded from fragmentation for MS2, dynamic exclusion for 30 s prevented repeated selection of precursors. Alternatively, samples were analyzed using an Ultimate 3000 RSLC nano (Thermo Fisher) coupled to an Orbitrap Exploris 480 mass spectrometer equipped with a FAIMS Pro interface for Field asymmetric ion mobility separation (Thermo Fisher). Peptides were pre-concentrated on an Acclaim PepMap 100 pre-column (75 µM x 2 cm, C18, 3 µM, 100 Å, Thermo Fisher) using the loading pump and buffer A** (water, 0.1% [v/v] TFA) with a flow of 7 µL/min for 5 min. Peptides were separated on 16-cm frit-less silica emitters (New Objective, 75-µm inner diameter), packed in-house with reversed-phase ReproSil-Pur C18 AQ 1.9 µm resin (Dr. Maisch, High Performance LC GmbH). Peptides were loaded on the column and eluted for 130 min using a segmented linear gradient of 5% to 95% solvent B (0 min: 5% B; 0–5 min: 5% B; 5–65 min: 20% B; 65–90 min: 35% B; 90–100 min: 55% B; 100–105 min: 95% B; 105–115 min: 95% B; 115–115.1 min: 5% B; 115.1–130 min: 5% B) (solvent A 0% ACN, 0.1% [v/v] FA; solvent B 80% [v/v] ACN, 0.1% [v/v] FA) at a flow rate of 300 nL/min. Mass spectra were acquired in data-dependent acquisition mode with a TOP_S method using a cycle time of 2 seconds. For field asymmetric ion mobility separation (FAIMS), two compensation voltages (−45 and −60) were applied, and the cycle time was set to 1 second for each experiment. MS spectra were acquired in the Orbitrap analyzer with a mass range of 320–1,200 m/z at a resolution of 60,000 FWHM and a normalized AGC target of 300%. Precursors were filtered using the MIPS option (MIPS mode = peptide), and the intensity threshold was set to 5,000. Precursors were selected with an isolation window of 1.6 m/z. HCD fragmentation was performed at a normalized collision energy of 30%. MS/MS spectra were acquired with a target value of 75% ions at a resolution of 15,000 FWHM, inject time set to auto and a fixed first mass of m/z 120. Peptides with a charge of +1, greater than 6, or with unassigned charge state were excluded from fragmentation for MS2.

Data analysis: Raw data were processed using MaxQuant software (version 1.6.3.4, http://www.maxquant.org/) (Cox & Mann, 2008) with label-free quantification (LFQ) and iBAQ enabled (Tyanova *et al*, 2016). MS/MS spectra were searched by the Andromeda search engine against a combined database containing the sequences from Arabidopsis (TAIR10_pep_20101214; ftp://ftp.arabidopsis.org/home/tair/Proteins/TAIR10_protein_lists/) and sequences of 248 common contaminant proteins and decoy sequences. Trypsin specificity was required and a maximum of two missed cleavages allowed. Minimal peptide length was set to seven amino acids. Carbamidomethylation of cysteine residues was set as fixed, oxidation of methionine and protein N-terminal acetylation as variable modifications. Peptide-spectrum-matches and proteins were retained if they were below a false discovery rate of 1%. Statistical analysis of the MaxLFQ values was carried out using Perseus (version 1.5.8.5, http://www.maxquant.org/). Quantified proteins were filtered for reverse hits and hits “identified by site”, MaxLFQ values were log2 transformed. After grouping samples by condition only those proteins were retained for the subsequent analysis that had two valid values in one of the conditions. Two-sample *t*-tests were performed using a permutation-based FDR of 5%. Alternatively, quantified proteins were grouped by condition and only those hits were retained that had 3 valid values in one of the conditions. Missing values were imputed from a normal distribution (1.8 downshift, separately for each column). The Perseus output was exported and further processed using Excel. Candidates with LFQ intensity value > 20 and log2 ratio > 2 compared to both WT and “YFP” AP-controls were retained (Supplemental Table S1).

### RNA extraction and reverse transcription quantitative PCR (RT-qPCR)

Total RNA from plant tissues was isolated using the Plant RNeasy kit (Qiagen), including DNase treatment, following the manufacturer’s protocol. cDNA synthesis was carried out using the Taqman Reverse Transcription Reagent kit (Thermo Fisher Scientific). Each RT-qPCR reaction contained cDNA equivalent to 3 ng of RNA, synthesized with a 1:1 mixture of oligo(dT) primers and random hexamers. RT-qPCR was performed using PowerUp SYBR Green Master Mix (Thermo Fisher Scientific) on a QuantStudio™ 5 Real-Time PCR System (Thermo Fisher Scientific). Gene expression levels were calculated using the ΔΔCT method (Livak & Schmittgen, 2001), with *PP2AA3* (*PROTEIN PHOSPHATASE 2A SUBUNIT A3*, AT1G13320) or *UBQ5* (*UBIQUITIN 5*, AT3G62250) as reference genes (Czechowski *et al*, 2005). Each experiment included three independent biological replicates.

### Transient protein expression, BiFC, and LCI assay in *Nicotiana benthamiana*

To transiently express proteins in *Nicotiana benthamiana,* constructs were introduced into Agrobacterium strain GV3101 and the OD values were monitored using a spectrophotometer, after which cultures were co-infiltrated alongside silencing inhibitor P19 (Lombardi *et al*, 2009) into young leaves of *N. benthamiana*. Plants were grown under LD conditions (16-h light/8-h dark) for 2 d. For the BiFC assay, the signal was visualized and documented using a confocal microscope (Leica SP8). For LCI assay, luciferin (1 mM) was sprayed evenly onto infiltrated leaves and kept in darkness for 7 mins, after which LUC activity was monitored with LB985 NightShade with indiGo software (Berthold Tech).

### Protein sequence analyses and structure modeling

Multiple sequence alignment of CAIR1 proteins was performed using MUSCLE in Jalview. CAIR1 structure was modeled using Alpha fold (Abramson *et al*, 2024). Three-dimensional structural models were visualized using PyMOL software.

### Catalase activity measurements

Catalase activity was measured as previously described (Aebi, 1984; Noctor *et al*, 2016). Briefly, 80 mg of 5-d-old seedlings was homogenized in 1.5 mL extraction buffer (100 mM PBS, pH 7.0) for 5 min at 4°C. The homogenate was centrifuged at 12,000 *g* for 15 min at 4°C, and the supernatant was collected. 100 μL of crude protein extract was diluted to a total volume of 3 mL reaction mixture in 100 mM PBS (pH 7.0) containing 30 mM H_2_O_2_. Catalase activity was measured spectrophotometrically at room temperature by monitoring the decrease in absorbance at 240 nm resulting from the decomposition of H_2_O_2_. Protein concentrations were determined using the Bio-Rad Protein Assay Dye Reagent Concentrate, following the manufacturer’s instructions.

### DAB staining

Reactive oxygen species (ROS) were detected in 10-d-old seedlings before and after a 24-h gas limitation treatment using 3,3′-diaminobenzidine (DAB) (Sigma-Aldrich) staining, following a previously described method (Daudi *et al*, 2012). Images were captured under a microscope.

### *F*v/*F*m measurement

The maximum quantum efficiency of PSII (*F*v/*F*m) was assayed by measuring plant chlorophyll fluorescence using a PSI (Photon Systems Instruments) FluorCam 800 MF, as previously described (Leonardelli *et al*, 2024). Before measurements, plates were incubated in darkness for 7 min which was confirmed to be sufficient to reach steady state under the plant growth conditions used (Murchie & Lawson, 2013). Maximum fluorescence in the dark (*F*m) was measured using saturating pulses of 2000 µmol m^-2^ s^-1^ (white light, 400–720 nm) for 960 ms, followed by orange-red light (620 nm) detection pulses (10 µs). *F*v/*F*m was calculated as *F*v/*F*m = (*F*m − *F*o)/*F*m, where *F*m is the maximal fluorescence and *F*o is the minimal fluorescence in the dark-adapted state (Baker, 2008).

### Hypocotyl length measurements

Seedlings were grown for 5 d under in white light (3.6 µmol m^-2^ s^-1^) supplemented with UV-B (1.5 µmol m^-2^ s^-1^) (+), or not (−) (Oravecz *et al*, 2006), after which approximately 30 seedlings per genotype or condition were aligned on an agar plate and scanned. The length of individual hypocotyls was quantified using ImageJ software.

### Extraction and quantification of anthocyanins

Anthocyanin content was quantified as previously described (Podolec *et al*, 2022). Briefly, seedlings were grown for 4 d in white light (3.6 µmol m^-2^ s^-1^) supplemented with UV-B (1.5 µmol m^-2^ s^-1^) (+), or not (−) (Oravecz *et al*, 2006), and approximately 20 mg of seedlings were collected for each genotype and condition. Samples were frozen in liquid nitrogen, ground to a fine powder, and extracted with 400 µL of extraction buffer (99% [v/v] methanol, 1% [v/v] HCl). Extraction was carried out for at least 1 h by incubating the samples on a rotary shaker at 4 °C. Following centrifugation for 5 min, the absorbance of 200 µL of the clarified supernatant was measured at 530 nm and 655 nm with a TECAN Spark 10M Multimode Microplate Reader. Relative anthocyanin content was calculated using the formula: (A530 – 0.25 * A655) / seedling mass (mg).

### Extraction of phenolic compounds and measurements of absorption spectra

Extraction of phenolic compounds and measurements of absorption spectra was performed as previously described (Leonardelli *et al*, 2024). In short, phenolic compounds were extracted from 50 mg of fresh plant material homogenized with a Silamat S6 (Ivodar Vivadent) in 400 μL of 80% (v/v) methanol. The homogenate was incubated for 10 minutes at 70°C with shaking at 800 *g*, followed by centrifugation at 12,000 *g* for 10 minutes at room temperature (RT). Subsequently, 200 µL of the clear supernatant was transferred to a transparent 96-well plate (UV-SATR MICROPLATE, Greiner), and absorbance spectra were recorded using a TECAN Spark 10M Multimode Microplate Reader.

### Yeast growth assay

The AD and BD constructs were co-transformed into the yeast strain Y2Hgold (Takara) and selected on SD/–Leu/–Trp double-dropout (DDO) plates. To assess the protein interactions, the transformed yeasts were placed on SD/–Leu/–Trp/–His triple-dropout (TDO) plates. The interactions were observed after 4 d at 30°C.

### Confocal laser-scanning microscopy

The localization of YFP- or GFP-tagged proteins was imaged using a Leica TCS SP8 confocal microscope (Leica, Bensheim, Germany). GFP and YFP were excited with Argon laser at 488 nm and the emitted fluorescence was detected between 493 and 550 nm.

### Statistical analysis

All data was analyzed and graphed with GraphPad Prism software (version 10.6.1).

## Acknowledgements

We would like to thank Lug Trémulot, Michael Hothorn, and Emilie Demarsy for helpful comments on the manuscript, Christophe Belin (University of Perpignan, France) and Dmitry Lapin (Max Planck Institute for Plant Breeding Research, Germany) for kindly providing seeds of the peroxisome maker line GFP-PTS1 and the Pro_35S_:StrepII-3xHA-YFP transgenic line, respectively, the European Arabidopsis Stock Centre (NASC) for providing various mutant seed material, Lothar Kalmbach (University of Neuchâtel, Switzerland) for plasmid pLOK180_pFR7m34GW, Richard Chappuis (University of Geneva) for transient transfection of HEK293 cells, Rubén Tenorio Berrío (University of Geneva) for help with GO term analysis, and Anne Harzen (Max-Planck Institute for Plant Breeding Research) for MS sample preparation. This work was supported by the Max Planck Society, the University of Geneva and the Swiss National Science Foundation (grant 310030_207716 to R.U.).

---

**Supplemental Figure S1.**
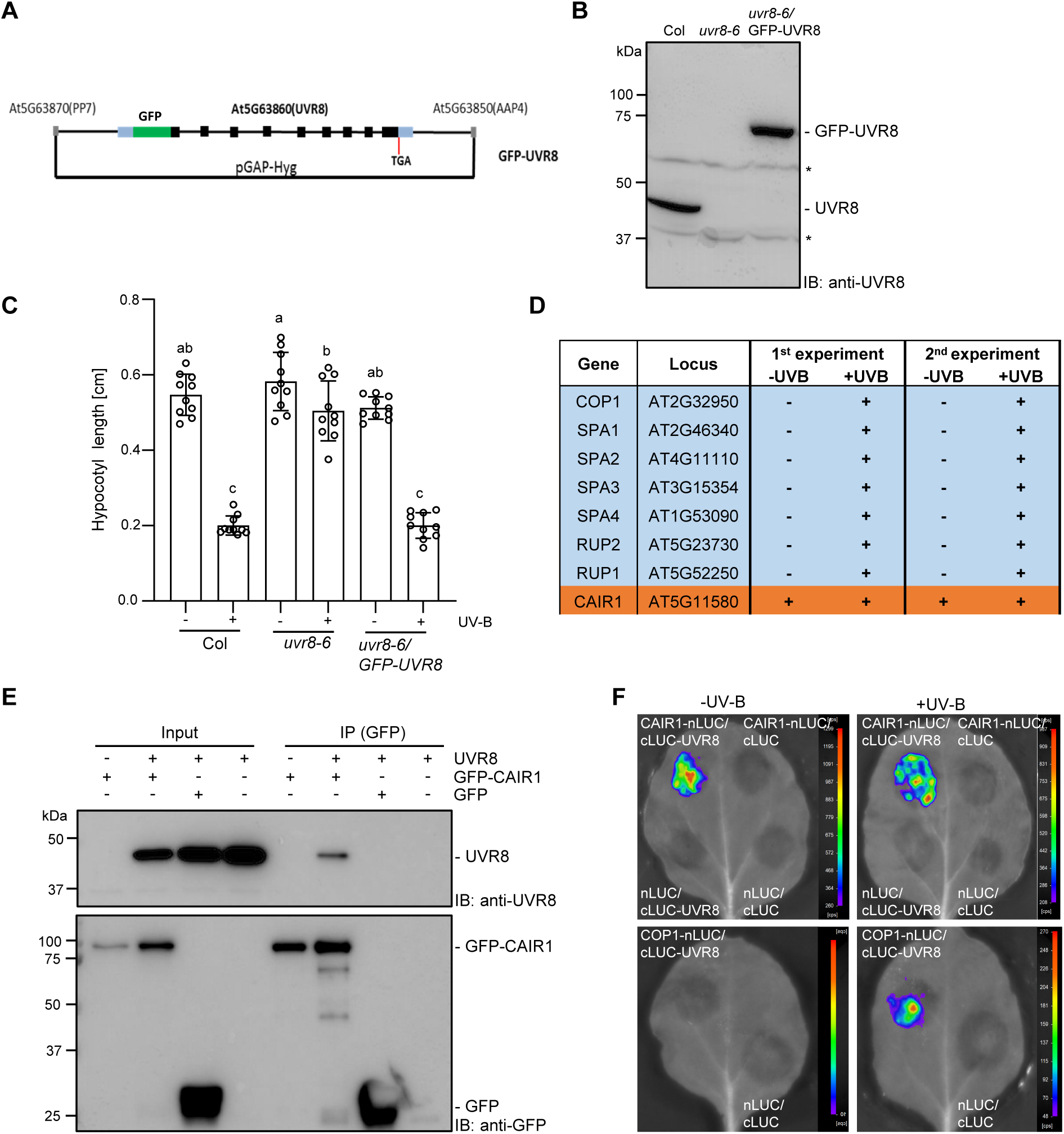
UVR8 interacts with a protein of unknown function (AT5G11580). **(A)** Schematic representation of the Pro_UVR8_:GFP-UVR8 genomic construct. Black boxes: exons of *UVR8*; black lines: introns, intergenic regions to neighboring genes At5g63870 (*PP7; SERINE/THREONINE PHOSPHATASE 7*) and At5G63850 (*AAP4’ AMINO ACID PERMEASE 4*), as well as pGAP-Hyg vector backbone; green box: GFP coding sequence; blue boxes: 5′ and 3′ UTRs of *UVR8*. Translational start ATG provided by inserted GFP sequence; *UVR8* TGA stop codon is indicated. **(B)** Immunoblot analysis of UVR8 and GFP-UVR8 levels in wild-type (Col), *uvr8-6*, and *uvr8-6*/Pro_UVR8_:GFP-UVR8 line #16 (*uvr8-6*/GFP-UVR8) seedlings. Asterisks (*) indicate cross-reacting unspecific bands. **(C)** Quantification of hypocotyl length of Col, *uvr8-6* and *uvr8-6*/GFP-UVR8 seedlings grown in white light (3.6 µmol m^-2^ s^-1^) supplemented (+) or not (-) with UV-B (1.5 µmol m^-2^ s^-1^). Individual seedling data points and means ± SD are shown (*n* = 10). Shared letters indicate no statistically significant difference between the means (P > 0.05), as determined by two-way ANOVAs followed by Tukey’s test for multiple comparisons. **(D)** Selected proteins identified in two independent AP–MS experiments of GFP-UVR8 complexes from plants grown in white light in the absence (-UV-B) or presence (+UV-B) of supplementary UV-B; “+” and “-” signs indicate, respectively, enrichment or not in the AP–MS analysis. Known UVR8 interactors and complex constituents are highlighted with a light blue background and the previously undescribed CAIR1 (AT5G11580) in orange. **(E)** Co-IP assay of endogenous UVR8 using GFP-trap purification, showing co-IP of UVR8 with GFP-CAIR1 but not GFP when co-expressed in HEK293 cells. IB, immunoblotting; IP, immunoprecipitation. **(F)** Transient expression in *N. benthamiana* leaves for LCI assay of the interaction of cLUC-UVR8 with co-expressed CAIR1-nLUC and COP1-nLUC (positive control in presence of UV-B), as well as the respective empty vector negative controls (nLUC and cLUC). Before imaging luciferase activity, the leaves were exposed to UV-B for 10 min (+UV-B), or not (-UV-B).

**Supplemental Figure S2.**
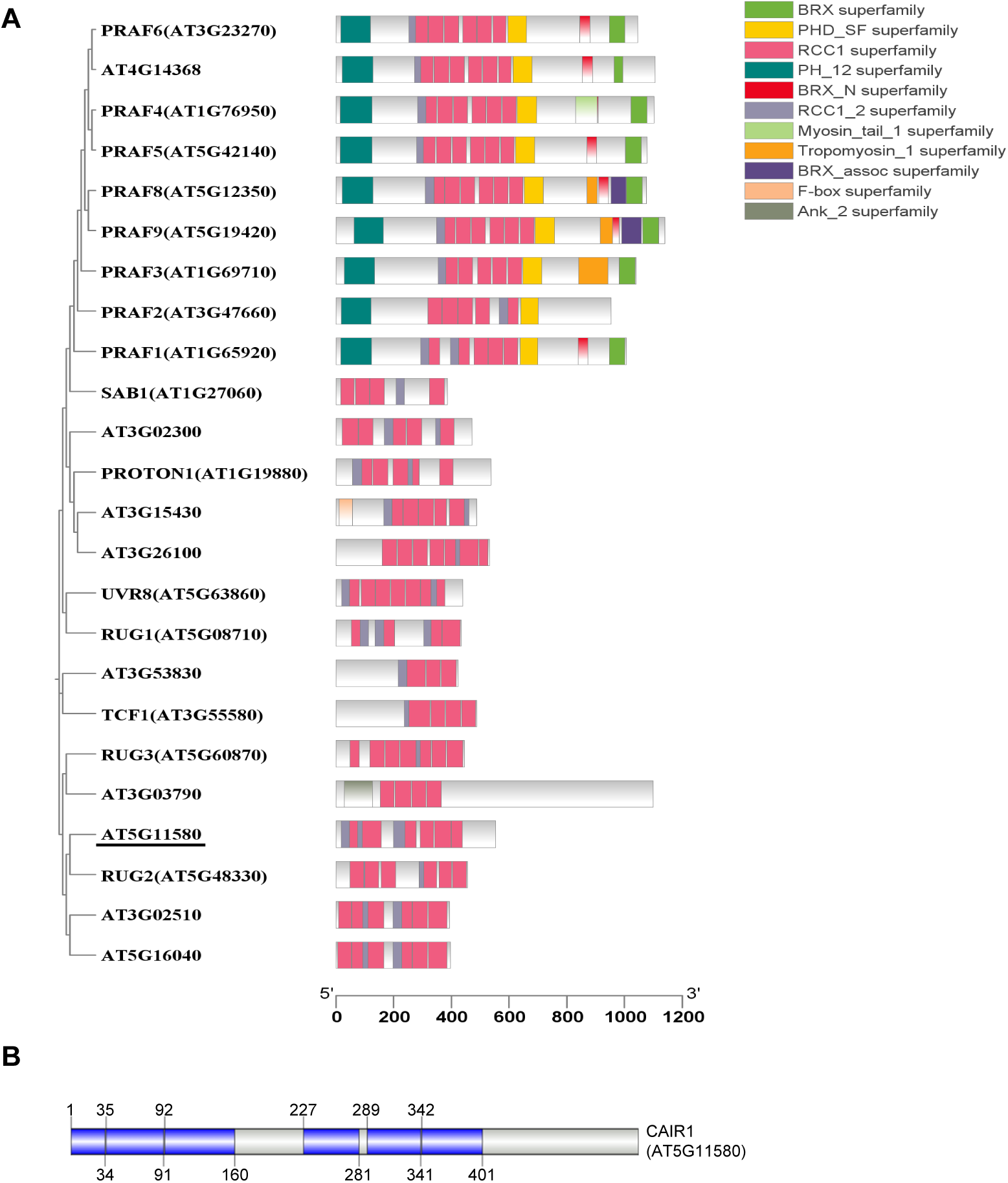
CAIR1 (AT5G11580) is a previously uncharacterized RCC1-like family member. **(A)** Phylogenetic tree and domain structures of Arabidopsis RCC1-like family members. The sequences of 24 RCC1 family proteins were downloaded from TAIR and aligned with Clustal Omega (https://www.ebi.ac.uk/jdispatcher/msa/clustalo). The phylogenetic tree was generated in MEGA-X using the default parameters. The domain architectures were retrieved from the Pfam protein families database (https://www.ncbi.nlm.nih.gov/Structure/bwrpsb/bwrpsb.cgi). Phylogenetic tree and structures were visualized schematically using TBtools-II ^121^. **(B)** CAIR1 features six RCC1 domains and an extended C-terminal region. The 553 amino acid–long CAIR1 sequence was obtained from TAIR. The domain was subsequently analyzed using the EBI database (https://www.ebi.ac.uk), and the results were visualized using IBS 2.0 (https://ibs.renlab.org/#/server). First (top) and last (bottom) amino acid residue for each RCC1 domain is given.

**Supplemental Figure S3.**
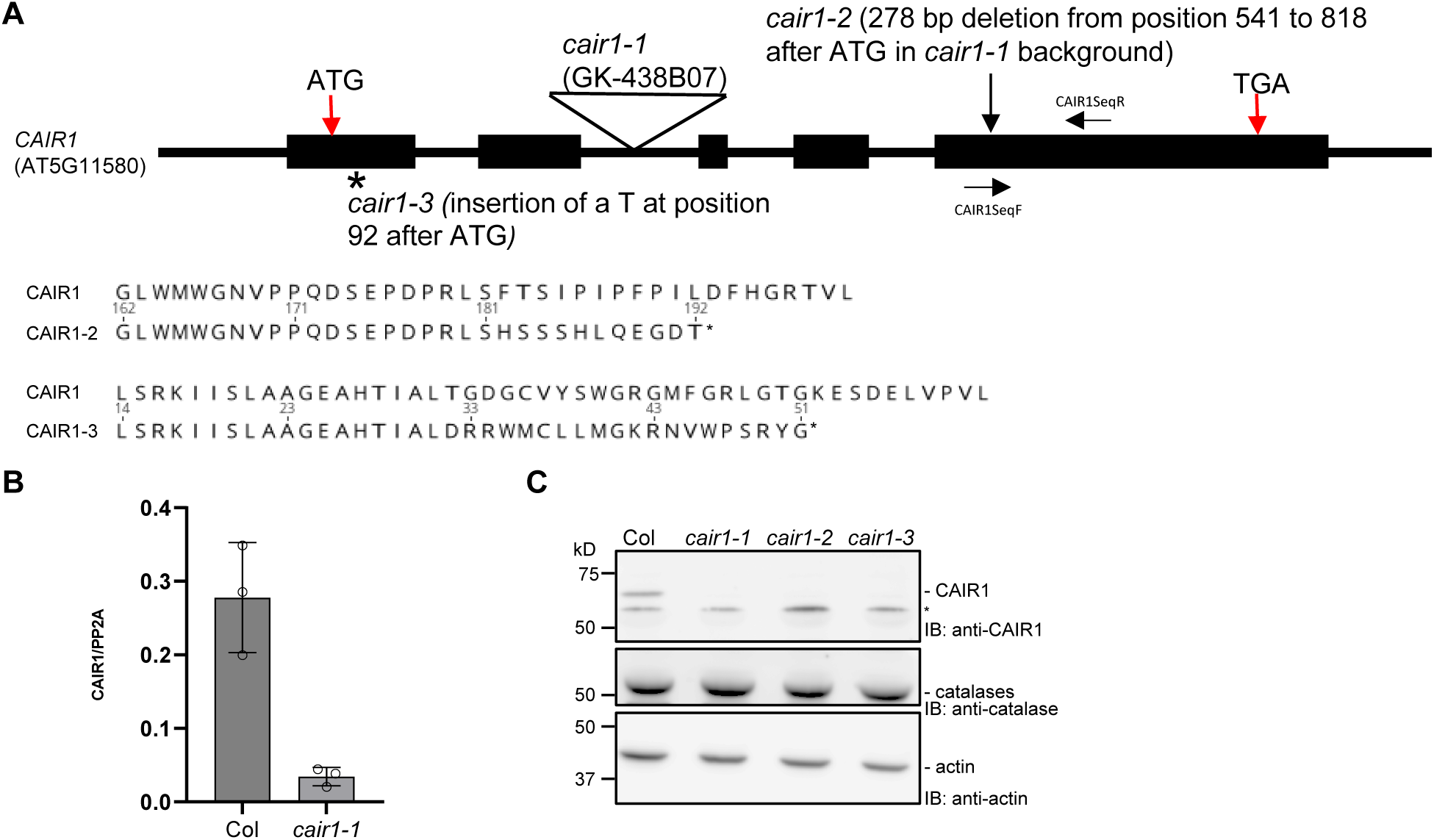
*cair1* mutant alleles. **(A)** Schematic structure of the *CAIR1* gene and mutant alleles. Black boxes indicate exons, black lines introns, as well as 5′ and 3′ UTRs. Translational start ATG and stop TGA codons are indicated with red arrows. Amino acid sequences and premature stop codons (*) for CRISPR/Cas9-generated *cair1-2* and *cair1-3* are indicated below adjacent to wild-type CAIR1 sequences. Approximate location of primers CAIR1SeqF and CAIR1SeqR used for RT-qPCR are indicated. **(B)** RT-qPCR analysis of *CAIR1* expression in wild type (Col) and *cair1-1* using primers CAIR1SeqF and CAIR1SeqR. Independent data points and means ± SD are shown (n = 3). **(C)** Immunoblot analysis of CAIR1 and catalase levels in 5-d-old *cair1-1*, *cair1-2*, and *cair1-3* seedlings compared to that in Col. *, unspecific band. Actin levels are shown as loading control.

**Supplemental Figure S4.**
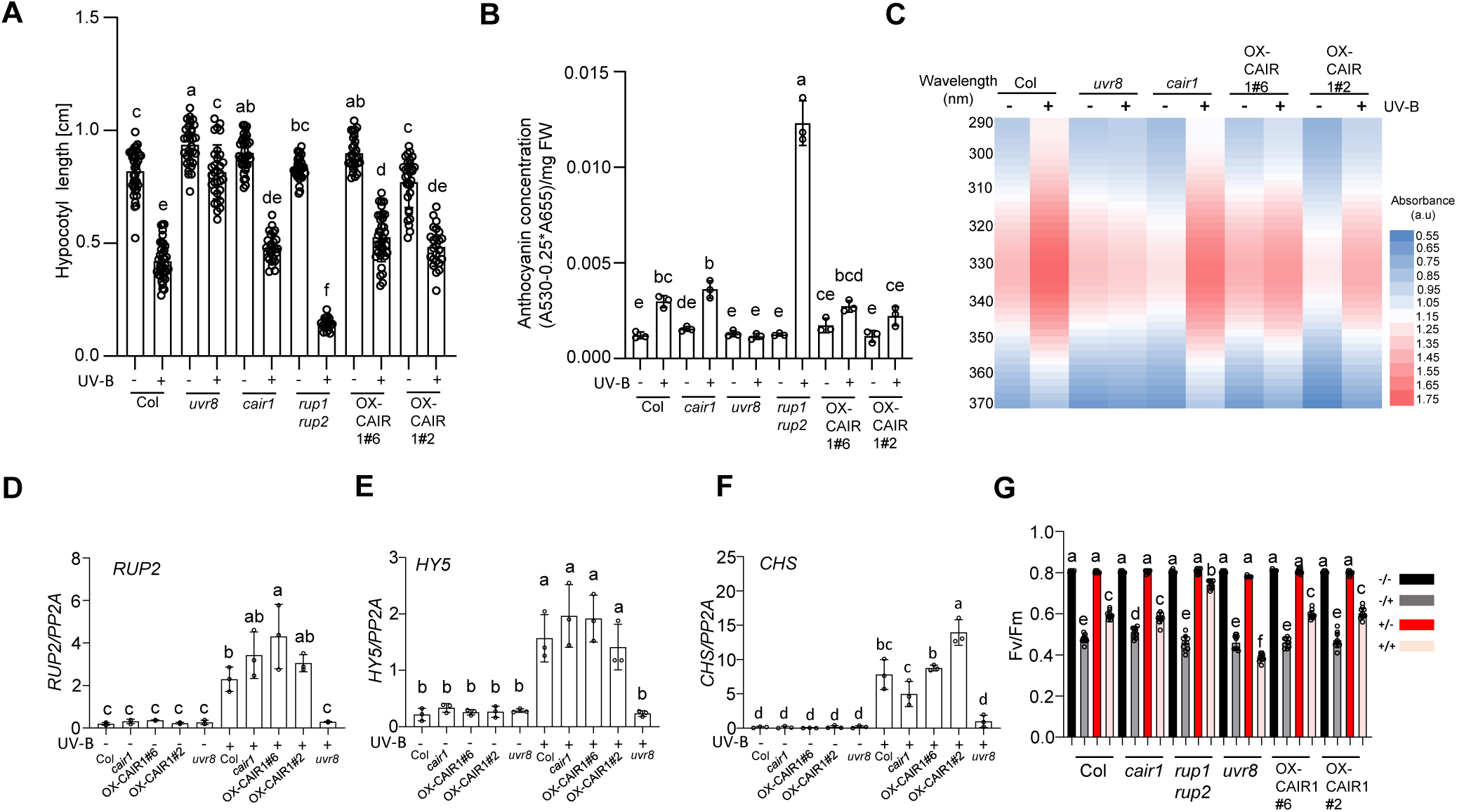
CAIR1 is not required for UV-B-induced photomorphogenic responses and UV-B acclimation. **(A–C)** Quantification of hypocotyl length (A), anthocyanin concentration (B), and absorbance spectra (280–400 nm) of methanolic extracts (C; UV-absorbing metabolites) of 5-d-old wild-type (Col), *uvr8-6* (*uvr8*), *rup1 rup2*, *cair1-1* (*cair1*), and two CAIR1 overexpression lines (OX-CAIR1 lines #2 and #6) seedlings grown in white light (3.6 µmol m^-2^ s^-1^) supplemented with UV-B (1.5 µmol m^-2^ s^-1^) (+), or not (−). (A) Individual seedling data points and means ± SD are shown (n > 20). (B) Triplicate individual data points and means ± SD are shown. **(D–F)** RT-qPCR analysis of *RUP2*, *HY5*, and *CHS* expression levels after UV-B treatment. Five-d-old Col, *uvr8*, *rup1 rup2*, *cair1*, and OX-CAIR1 lines #2 and #6 seedlings were grown in white light (3.6 µmol m^-2^ s^-1^), then exposed (+) or not (-) to 3 h of narrowband UV-B (1.5 µmol m^-2^ s^-1^). Triplicate independent experiment data points and means ± SD are shown. **(G)** *F*v/*F*m measurements of Col, *uvr8*, *rup1 rup2*, *cair1*, and OX-CAIR1 lines #2 and #6 seedlings. Seven-d-old light-grown seedlings were exposed (acclimated, +/−) or not (−/−) to UV-B for 3 d with supplementary narrowband UV-B (0.08 mW cm^−2^). After 10 d, acclimated and non-acclimated plants were exposed to 2 h of broadband UV stress irradiation (non-acclimated and stressed −/+; acclimated and stressed +/+; 2.2 mW cm^−2^). Independent plant data points and means ± SD are shown (n = 10). (A–G) Shared letters indicate no statistically significant difference between the means (P > 0.05), as determined by two-way ANOVAs followed by Tukey’s test for multiple comparisons.

**Supplemental Figure S5.**
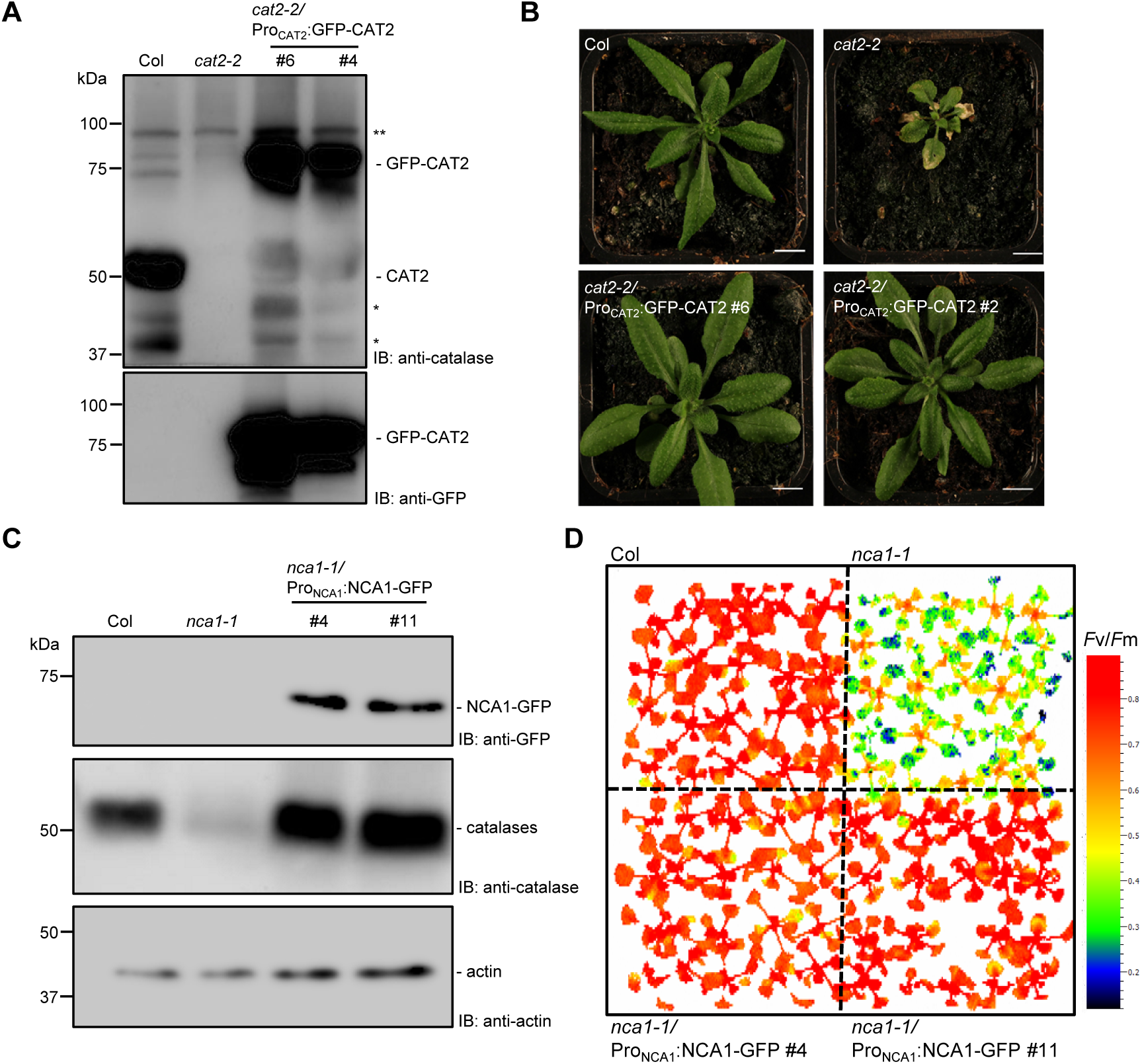
Functional complementation of *cat2-2* and *nca1-1* mutant phenotypes by Pro_CAT2_:GFP-CAT2 and Pro_NCA1_:NCA1-GFP genomic clones, respectively. **(A)** Immunoblot analysis of 3-week-old *cat2-2*/Pro_CAT2_:GFP-CAT2 complementation lines #6 and #4 compared to wild type (Col) and *cat2-2* using anti-catalase and anti-GFP antibodies. IB, immunoblotting. *, degradation products; **, unspecific cross-reacting band. **(B)** The transgenic lines expressing GFP-CAT2 fully rescue the small size and leaf lesion phenotypes of *cat2-2* under long-day (LD) conditions. **(C, D)** The transgenic lines expressing NCA1-GFP fully restore catalase protein stability and alleviate the oxidative stress sensitivity of *nca1-1*. (C) Immunoblot analysis of catalase and NCA1-GFP levels in 7-d-old *nca1-1*/Pro_NCA1_:GFP-CAT2 complementation lines #4 and #11 compared to Col and *nca1-1*. Actin levels are shown as loading control. IB, immunoblotting. (D) *F*v/*F*m measurements of 7-d-old seedlings of the indicated genotypes grown for 2 d under gas limitation (enhanced photorespiration).

**Supplemental Figure S6.**
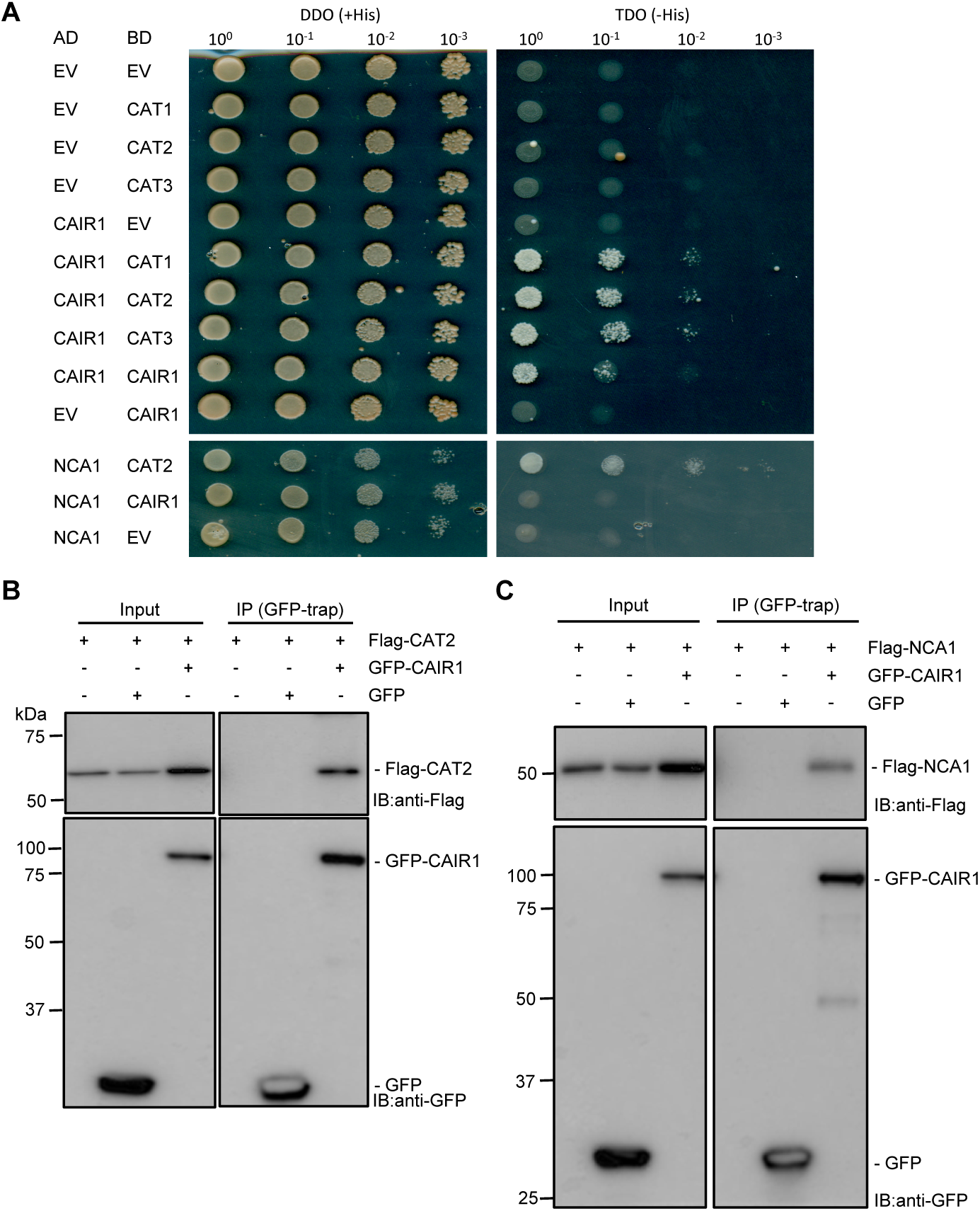
CAIR1 interacts with catalases and NCA1 in yeast and HEK293 cells. **(A)** Yeast two-hybrid analysis of CAIR1, catalases CAT1, CAT2, CAT3, and NCA1. Tenfold serial dilutions of transformed yeast spotted on SD/−Trp/−Leu (DDO; nonselective for interaction) and SD/−Trp/−Leu/−His (TDO; selective) plates. AD, activation domain; BD, DNA binding domain; EV, empty vector. **(B, C)** HEK293T cells were co-transfected to express the indicated proteins, and GFP-CAIR1 and GFP (negative control) were immunoprecipitated using GFP-trap nanobodies. The IP signals (GFP-CAIR1 and GFP) and the co-IP signals (Flag-CAT2 and Flag-NCA1) were detected by immunoblots probed with antibodies against GFP- and Flag-epitopes, respectively. IP, immunoprecipitation; IB: immunoblotting.

**Supplemental Figure S7.**
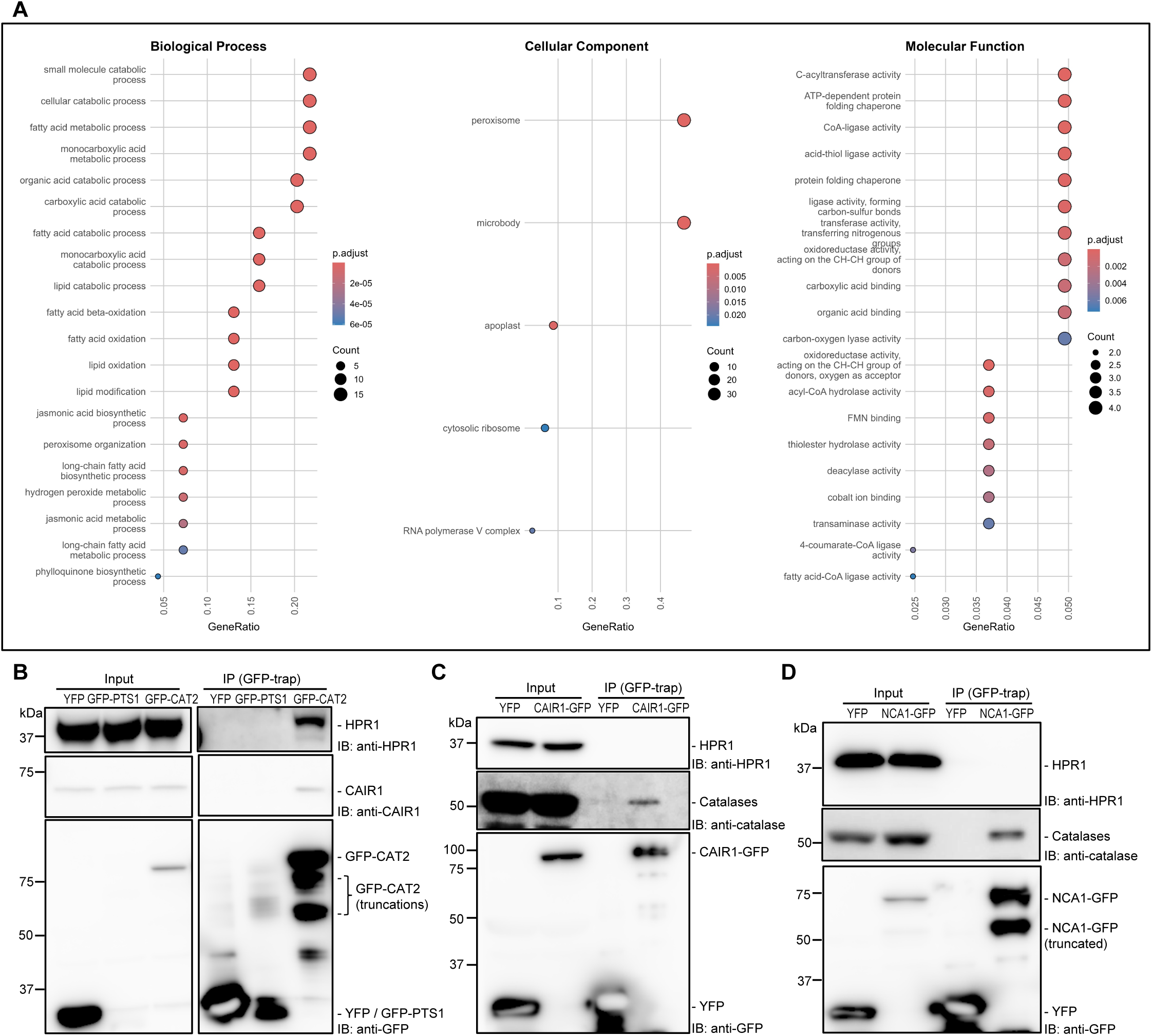
The interactome of CAT2 in Arabidopsis. **(A)** Enriched GO terms in Biological Process, Cellular Component, and Molecular Function among GFP-CAT2 interaction candidates (Supplemental Table S1). **(B)** Co-IP of endogenous HPR1 and CAIR1 using GFP-trap purification from extracts of 15-d-old *cat2-2*/Pro_CAT2_:GFP-CAT2 (GFP-CAT2) seedlings. **(C)** Co-IP of endogenous catalases but not HPR1 using GFP-trap purification from extracts of 15-d-old *cair1-1*/Pro_CAIR1_:CAIR1-GFP (CAIR1-GFP) seedlings. **(D)** Co-IP of endogenous catalases but not HPR1 using GFP-trap purification from extracts of 15-d-old *nca1-1*/Pro_NCA1_:NCA1-GFP (NCA1-GFP) seedlings. (A–C) Pro_35S_:StrepII-3xHA-YFP (YFP) and/or Pro_35S_:GFP-PTS1 (GFP-PTS1) lines were used as negative controls. IB, immunoblotting; IP, immunoprecipitation.

**Supplemental Figure S8.**
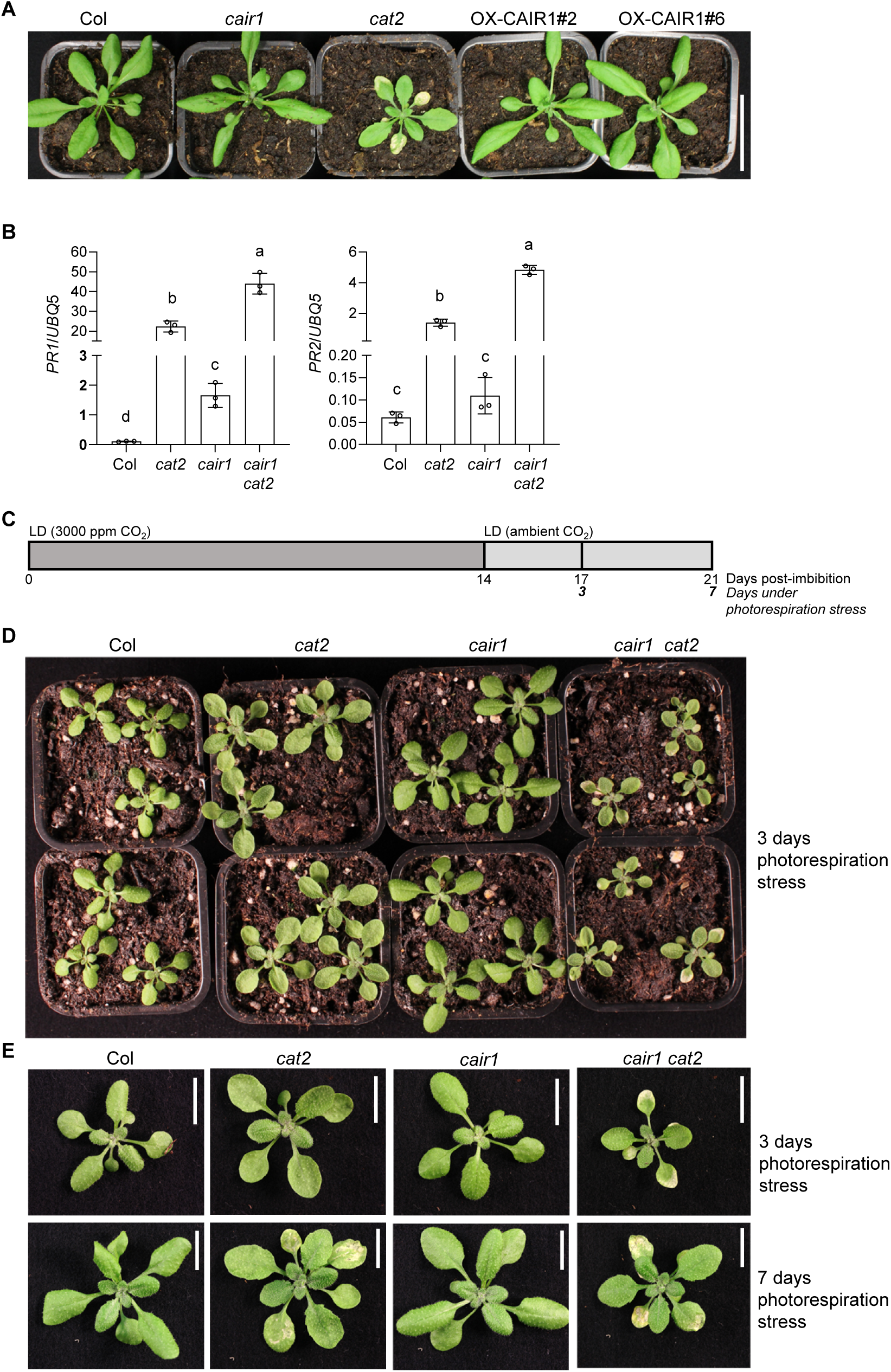
*cair1* enhances *cat2* leaf lesion phenotype under acute photorespiration stress in LD. **(A)** Photographs of 15-d-old wild-type (Col), *cair1-1* (*cair1*), *cat2-2* (*cat2*), and two CAIR1 overexpression lines (OX-CAIR1 lines #2 and #6) plants grown on soil under long-day (LD) conditions. Scale bar: 3 cm. **(B)** RT-qPCR analysis of *PR1* and *PR2* gene expression in 15-d-old wild-type (Col), *cair1*, *cat2*, and *cair1 cat2* plants grown on soil under long-day (LD) conditions. Independent experiment data points and means ± SD (n = 3) are shown. **(C)** Scheme for acute photorespiration stress treatment. Plants were grown for two weeks in LD conditions under 3,000 ppm CO_2_ and then transferred to a standard LD chamber with ambient CO_2_ levels. **(D, E)** Photographs of Col, *cat2*, *cair1*, and *cair1-1 cat2-2* (*cair1 cat2*) plants 3 and 7 d after the transfer of 14-d-old plants to acute photorespiration stress as outlined in panel B. (C) Overview of plants 3 d after photorespiration stress. (D) Leaf lesions on individual plants on black background. Scale bars: 1 cm.

**Supplemental Figure S9.**
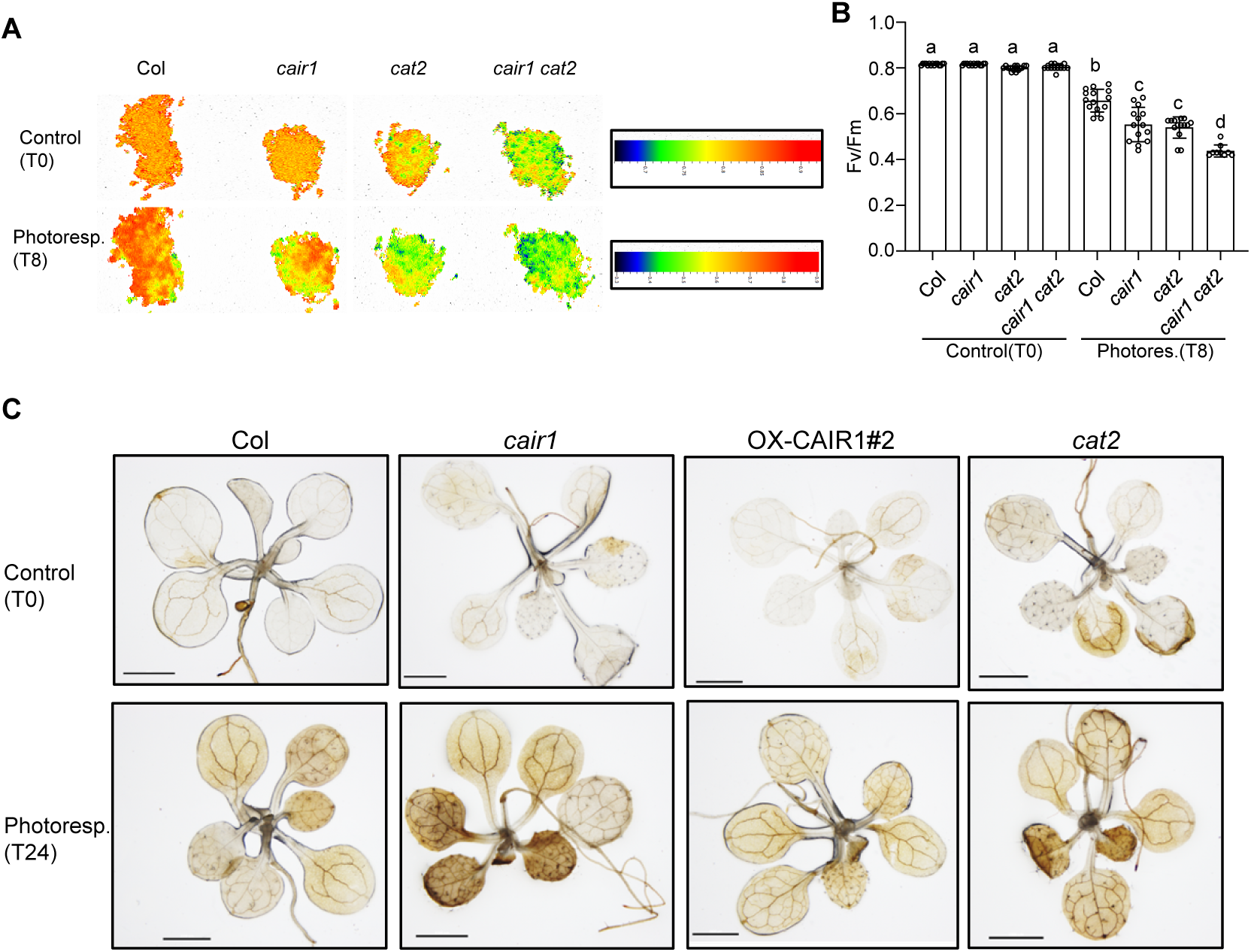
*cair1* exhibits sensitivity to oxidative stress. **(A)** Response to photorespiration–promoting growth conditions. False-color images representing *F*v/*F*m values of 10-d-old wild-type (Col), *cair1-1* (*cair1*), *cat2-2* (*cat2*), and *cair1-1 cat2-2* (*cair1 cat2*) seedlings before and after 8-h exposure to photorespiration-promoting growth conditions. Triplicate replicate experiments were performed, and a representative plate is shown. **(B)** Quantitative *F*v/*F*m measurements of (A). Independent plant data points and means ± SD are shown. Shared letters indicate no statistically significant difference between the means (P > 0.05), as determined by two-way ANOVAs followed by Tukey’s test for multiple comparisons. **(C)** DAB staining for H_2_O_2_ in 10-d-old Col, *cair1*, *cat2*, and CAIR1 overexpression line (OX-CAIR1#2) seedlings before and after 24-h exposure to photorespiration–promoting growth conditions. Scale bars = 200 μm.

**Supplemental Figure S10.**
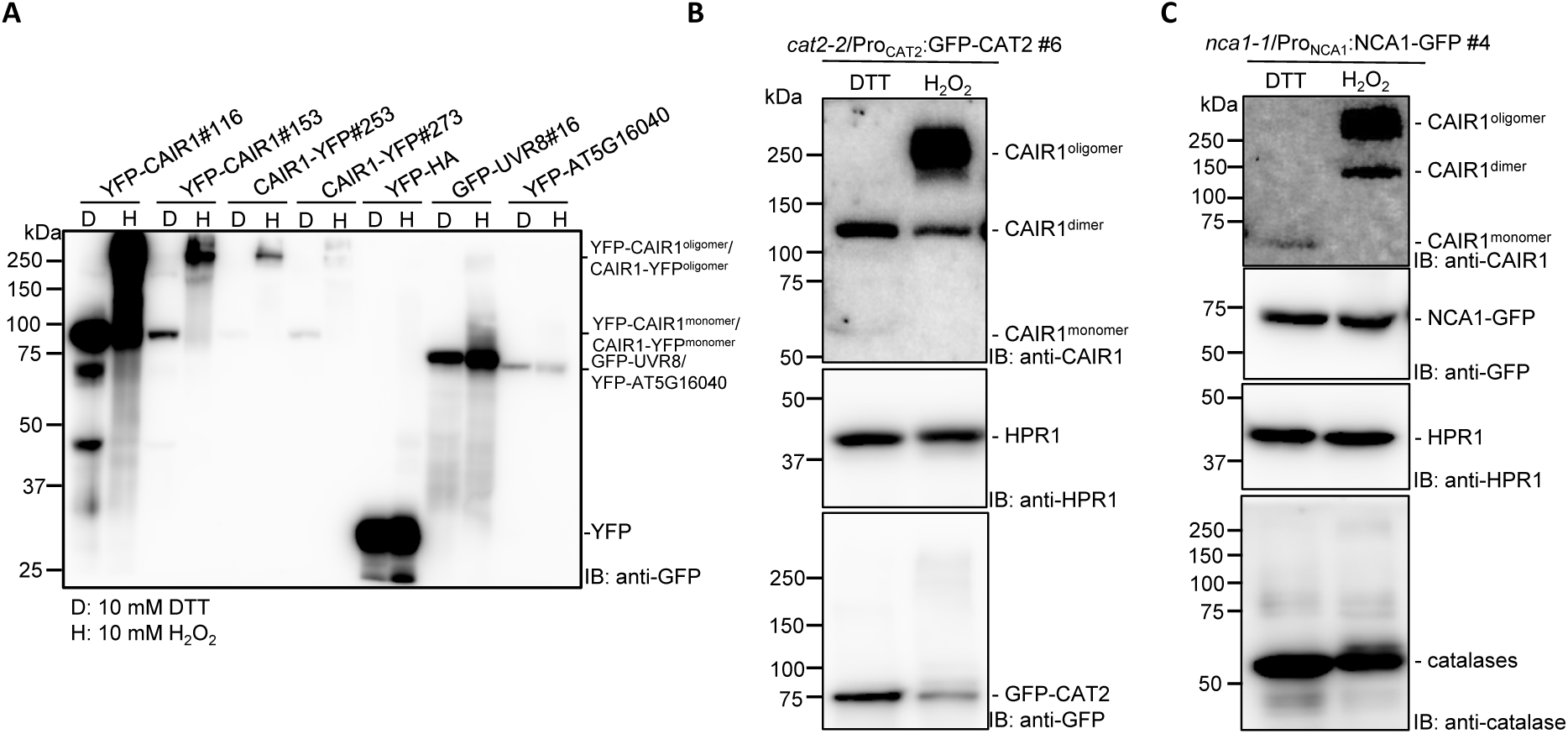
Specificity of redox–mediated CAIR1 oligomerization. **(A)** YFP-CAIR1 shows oligomerization under oxidative conditions (10 mM H_2_O_2_), unlike the other tested YFP- or GFP-tagged proteins. Proteins were extracted from 5-d-old *cair1-1*/Pro_35S_:YFP-CAIR1 #116 and #153, *cair1-1*/Pro_35S_:CAIR1-YFP #253 and #273, Col/Pro_35S_:YFP-HA, *uvr8-6*/Pro_UVR8_:GFP-UVR8 #16, Col/Pro_35S_:YFP-AT5G16040 seedlings. Then, the protein extracts were divided into two tubes to which 10 mM DTT (D) or 10 mM H_2_O_2_ (H) was added, respectively. **(B, C)** In contrast to CAIR1, no redox-dependent oligomerization of HPR1, GFP-CAT2, NCA1-GFP, or endogenous catalases was detected. Protein extracts from 5-d-old *cat2-2*/Pro_CAT2_:GFP-CAT2 #6 and *nca1-1*/Pro_NCA1_:NCA1-GFP #4 seedlings were treated with 10 mM DTT or 10 mM H_2_O_2_.

**Supplemental Figure S11.**
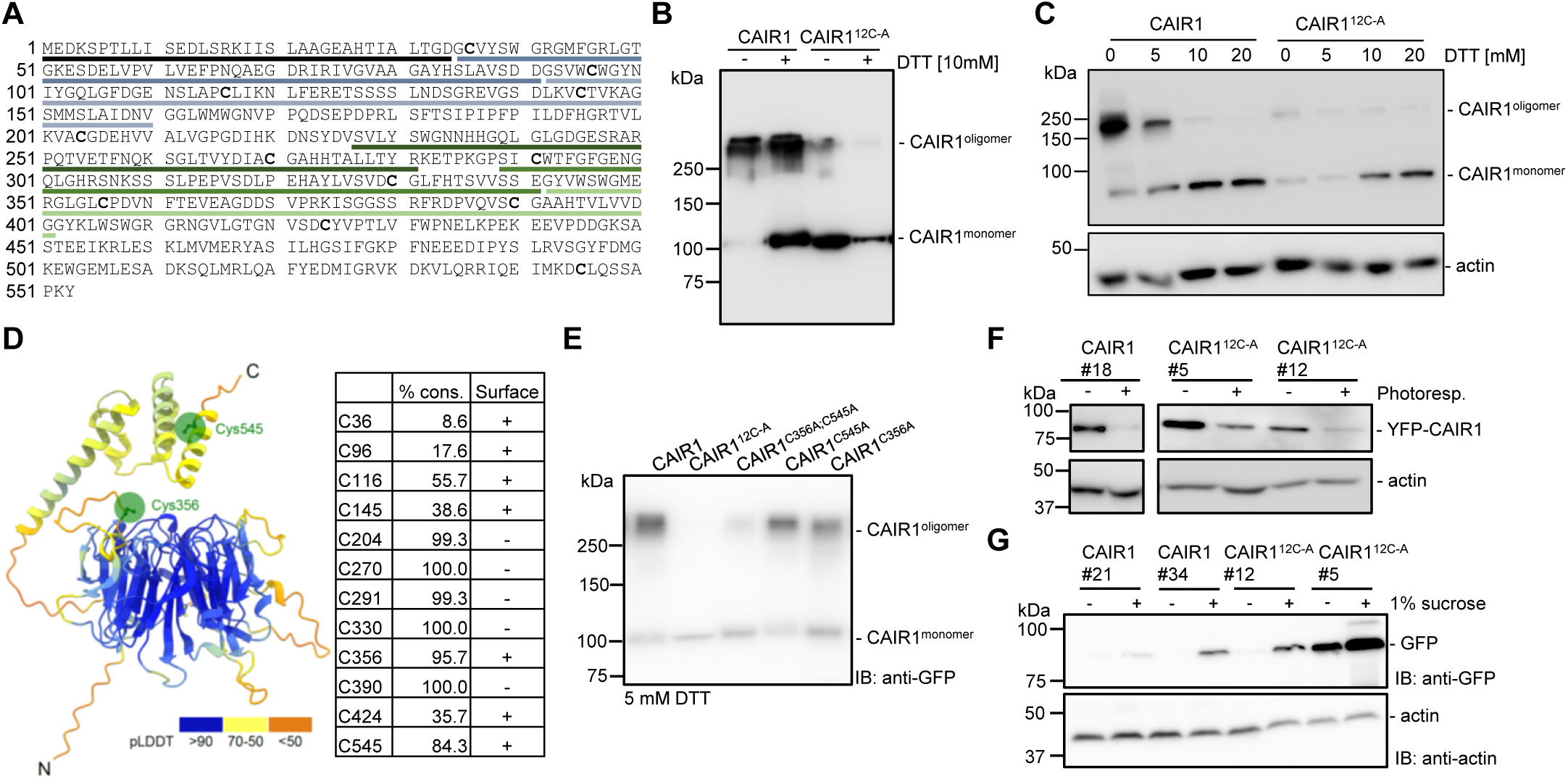
Redox-regulated oligomerization of CAIR1 depends on cysteines 356 and 545. **(A)** CAIR1 contains 12 cysteine residues highlighted in bold. The six RLDs are underlined in different colors (see also Supplemental Figure S2B). **(B)** YFP-CAIR1 (CAIR1) and YFP-CAIR1^12C-A^ (CAIR1^12C-A^) were transiently expressed in *N. benthamiana* and their oligomeric state in protein extracts in the presence of absence of 10 mM DTT was analyzed by anti-GFP immunoblot. **(C)** Proteins were extracted from 5-d-old *cair1-1*/Pro_35S_:YFP-CAIR1 #33 (CAIR1) and *cair1-1*/Pro_35S_:YFP-CAIR1^12C-A^ #25 (CAIR1^12C-A^) seedlings and CAIR1 oligomeric state in protein extracts containing different concentrations of DTT was analyzed by anti-GFP immunoblot. Actin levels are shown as loading control. **(D)** A ribbon diagram of the AlphaFold3 model of CAIR1 with surface-exposed cysteine 356 and 545 residues highlighted, alongside the % conservation of the 12 cysteine residues in plants. **(E)** YFP-CAIR1 variants were transiently expressed in *N. benthamiana* and their oligomeric state in protein extracts in the presence of 5 mM DTT was analyzed by anti-GFP immunoblot. **(F)** Mutation of the cysteine residues in CAIR1 did not alter ROS-mediated degradation of the protein. Six-d-old *cair1-1*/Pro_35S_:YFP-CAIR1 #18, *cair1-1*/Pro_35S_:YFP-CAIR1^12C-A^ #12, and *cair1-1*/Pro35S:YFP-CAIR1^12C-A^ #5 seedlings were transferred from a high CO_2_ chamber to conditions promoting enhanced photorespiration for 4 d. **(G)** The cysteine residues in CAIR1 were not essential for its sucrose-dependent stability. Total proteins were extracted from 6-d-old *cair1-1*/Pro_35S_:YFP-CAIR1 #21, *cair1-1*/Pro_35S_:YFP-CAIR1 #34, *cair1-1*/Pro_35S_:YFP-CAIR1^12C-A^ #12, and *cair1-1*/Pro_35S_:YFP-CAIR1^12C-A^ #5 seedlings grown in the absence (-) or presence of 1% sucrose (+) in the growth medium.

**Supplemental Figure S12.**
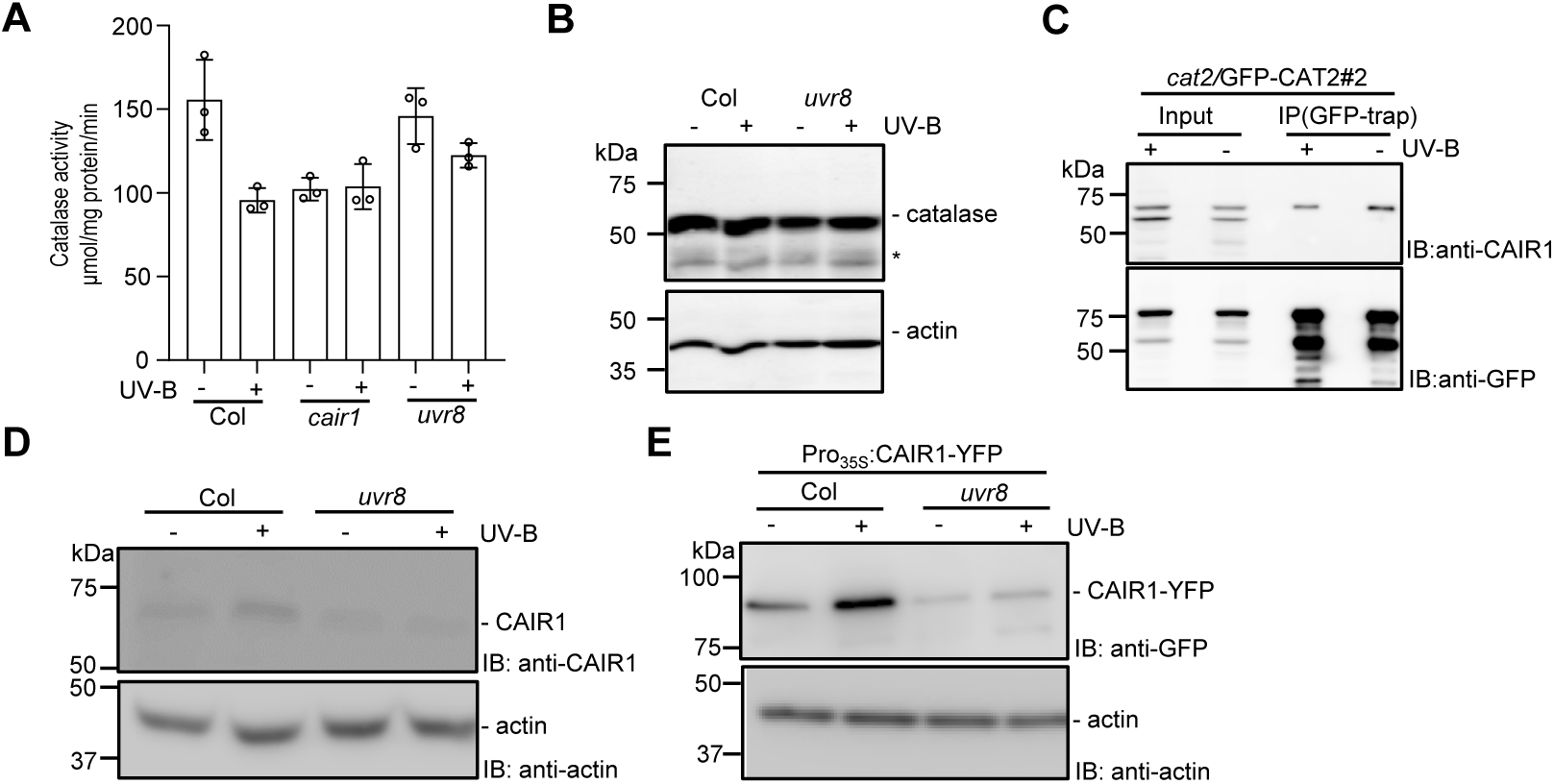
UV-B inhibits catalase activity through decreasing interaction of CAIR1 and catalase. **(A)** Catalase activity was measured in protein extracts from 5-d-old wild-type (Col), *cair1-1* (*cair1*), and *uvr8-6* (*uvr8*) seedlings. Data shown are means ± SD of 3 independent experiments. Shared letters indicate no statistically significant difference between the means (P > 0.05), as determined by two-way ANOVAs followed by Tukey’s test for multiple comparisons. **(B)** Immunoblot analysis of catalase and actin (loading control) protein levels in 5-d-old Col and *uvr8-6* seedlings grown with or without supplementary UV-B. Asterisk (*) indicates an unspecific band. **(C)** UV-B decreases interaction of catalase and CAIR1 in planta. Proteins were extracted from 5-d-old *cat2-2*/Pro_CAT2_:GFP-CAT2 #2 (*cat2*/GFP-CAT2#2) seedlings. **(D, E)** UV-B stabilizes CAIR1 in a UVR8-dependent manner. Proteins were extracted from 5-d-old (D) Col, *uvr8-6*, and (E) Col/Pro_35S_:CAIR1-YFP and *uvr8-6*/Pro_35S_:CAIR1-YFP seedlings. (A–E) Seedlings were grown in white light (3.6 µmol m^-2^ s^-1^) supplemented with UV-B (0.06 mW cm^-2^) (+), or not (−).

## Notes

### Competing Interest Statement

The authors have declared no competing interest.

